# The gonococcal vaccine candidate antigen NGO1701 is a *N. gonorrhoeae* periplasmic copper storage protein

**DOI:** 10.1101/2025.05.01.651437

**Authors:** Shea K. Roe, Rafał Mazgaj, Tianmou Zhu, Mariam Esmaeeli, Lisa A. Lewis, Caroline Genco, Kevin J. Waldron, Paola Massari

## Abstract

The increasing worldwide trend of antibiotic-resistant *Neisseria gonorrhoeae* strains highlights the urgent need for new therapeutic strategies against this sexually transmitted pathogen, including a gonococcal vaccine. We previously designed a bioinformatics-based candidate selection pipeline (CASS) and identified potential novel gonococcal vaccine targets among hypothetical proteins expressed during natural human infection. One of these candidates, NGO1701, is a predicted periplasmic four-helix bundle protein with amino acid sequence homology to the copper storage protein 1 (Csp1) from *Methylosinus trichosporium* OB3b. In this study, we confirmed that purified NGO1701 binds 15 Cu(I) ions per monomer *in vitro*, supporting its function as Csp in *N. gonorrhoeae*. Using a *ngo1701* deletion mutant generated in *N. gonorrhoeae* F62, we investigated its role in bacteria physiology. We showed that ablation of Csp was not limiting for bacterial growth and fitness *in vitro*, but the Δ*csp* strain became significantly more susceptible to copper mediated toxicity. This phenotype was rescued by *csp* gene complementation, indicating a role in protection against copper toxicity. Our results indicate that Csp participates in periplasmic copper homeostasis in *N. gonorrhoeae,* buffering excess copper to reduce toxicity and playing a putative role in copper delivery to important copper-enzymes. Csp does not appear to be involved in bacterial host cell interaction and activation *in vitro*, since no difference in the ability of *N. gonorrhoeae* to adhere/invade epithelial cells or induce IL-8 secretion was reported among wild type, *csp* deletion mutant and complemented strains. Furthermore, sera from mice immunized with NGO1701 failed to recognize Δ*csp* by dot blot and ELISA, and the sera’s ability to kill *N. gonorrhoeae* was abrogated against Δ*csp*. However, both functions were restored after gene complementation, supporting the relevance of Csp as a potential vaccine target. Allelic analysis of Neisseria species revealed that this gene is absent in *N. meningitidis*, thus making it a gonococcal-specific target.

**Author Summary:** Copper is essential for bacterial metabolism but can be toxic in excess. Here, we identify NGO1701 as a copper storage protein (Csp) in *Neisseria gonorrhoeae*, capable of sequestering Cu(I) ions. Deletion of *csp* led to increased copper sensitivity, while overexpression restored resistance, suggesting a role in copper homeostasis. The Δ*csp* mutant also showed reduced growth in cobalt and manganese, likely due to metal interference by copper toxicity. Beyond detoxification, Csp may supply copper to essential cuproenzymes like cytochrome *cbb*3 oxidase and nitric oxide reductase, which support bacterial survival under host-imposed stress. Although Csp is not required for *N. gonorrhoeae* host cell interactions, it is a strong immune target. Immune recognition of *N. gonorrhoeae* Δ*csp* by anti- NGO1701 mouse sera was nearly abolished and the serum bactericidal activity was abrogated compared to *N. gonorrhoeae* F62 wild type bacteria, highlighting Csp’s potential as a target for therapeutic or vaccine strategies against *N. gonorrhoeae*.

## Introduction

*Neisseria gonorrhoeae* is a gram-negative bacterium and the etiological agent of gonorrhea, a human sexually transmitted disease (STD). The incidence of gonorrhea has increased substantially over the past decade, with approximately 82.6 million cases reported annually worldwide and over 600,000 cases occurring in the United States [1]. While in men gonococcal infections are predominantly symptomatic, facilitating diagnosis and treatment, women are often asymptomatic or present with nonspecific symptoms. This can delay diagnosis and treatment, and lead to severe reproductive tract sequelae, such as endometritis, pelvic inflammatory disease (PID), ectopic pregnancy, and infertility. Disseminated gonococcal infections (DGI) and co-infections with *Chlamydia trachomatis*, *Treponema pallidum* and HIV are also well-documented [1]. Rising development of antimicrobial resistance has significantly complicated treatment of gonorrhea; fluoroquinolone resistance and rising levels of cefixime mean inhibitory concentrations (MICs) have excluded these antibiotics from treatment guidelines in the United States. Ceftriaxone is currently the sole first-line treatment suggested by the CDC, but resistance to this antibiotic has already been reported outside the U.S. [2–4]. The emergence and global spread of multidrug-resistant *N. gonorrhoeae* strains is a stark warning of the possibility of untreatable gonorrhea making new therapeutic options essential. A safe and effective vaccine would be the ideal solution.

Although both gonococcal and meningococcal outer membrane vesicles are being currently evaluated for efficacy against gonorrhea, gonococcal antigens continue to be explored [5], with the most advanced being the lipooligosaccharide (LOS) epitope recognized by the 2C7 Mab (and thus referred to as 2C7 epitope) [6, 7]. Preclinical research based on reverse vaccinology, transcriptomics and bioinformatics has led to discovery of conserved and surface-exposed antigens. Our group has developed a Candidate Antigen Selection Strategy (CASS) pipeline designed by integrating gonococcal protein expression in mucosal samples from men and women naturally infected with *N. gonorrhoeae* with predictions of protein surface exposure, immunogenicity, conservation and structure features [8–10]. With this approach, we identified several gonococcal hypothetical proteins as potential new vaccine targets. Three of these candidates (NGO0690, NGO0948 and NGO1701) were tested in mice and showed induction of antibodies with bactericidal activity, were recognized by IgG antibodies in sera from men and women naturally infected with *N. gonorrhoeae*, and where evaluated as a multi-antigen vaccine in mice [8, 11].

NGO1701 is a homologue of Csp1, a member of the family of copper storage proteins (Csps) characterized in the methanotrophic bacterium *Methylosinus trichosporium* OB3b [12]. Genes encoding cytosolic Csp homologues are found in diverse bacterial genomes, including pathogenic bacteria [13], while Csps that possess the twin arginine translocase (TAT) targeting sequence for secretion out of the cytoplasm are rare outside of the methanotrophic bacteria. Csps adopt a four-helix bundle fold that forms a tube structure lined with Cys residues that coordinate Cu(I) ions for copper storage and supply to copper-requiring enzymes. However, the influence of copper-buffering by cytosolic Csps in bacterial resistance to copper toxicity remains unclear, and a role for secreted Csps in copper resistance is yet untested. Crucially, no function of a Csp protein has been demonstrated in any pathogenic bacterium, despite copper homeostasis being essential to pathogens, and copper toxicity being a known weapon in the nutritional immunity arsenal of immune cells used to fight invading pathogens [14].

In this study, we investigated the function of NGO1701 in *N. gonorrhoeae,* which we here designate as the gonococcal Csp. Consistent with its homology to characterized Csp homologues, we demonstrated its ability to bind large quantities of copper *in vitro*. By generating a *csp* deletion mutant strain, we showed its role in *N. gonorrhoeae* resistance to copper toxicity and in periplasmic copper homeostasis. Together, our studies define an important function for Csp in *N. gonorrhoeae* and strengthen the evidence base for this protein as a target antigen for future anti-gonorrhea vaccine development.

## Results

### Sequence analysis and putative function of NGO1701

We previously reported that NGO1701 (WP_003689877.1) has sequence homology to a TAT_Cys_rich four helix bundle copper-binding protein of the DUF326 superfamily [8]. It was previously shown that a similar protein, Csp1 (copper storage protein 1) is expressed by the methanotrophic bacterium *M. trichosporium* OB3b bacteria [12], which functions in storage of Cu(I) ions for supply to its copper-dependent methane monooxygenase enzyme [12]. A sequence alignment of NGO1701 with Csp1 (*Mt*Csp1) and its paralogue Csp2 (*Mt*Csp2) from *M. trichosporium* revealed that they share 38% and 34% sequence identity with the *N. gonorrhoeae* homologue, respectively (**Fig 1**). Crucially, all the Cys residues shown to coordinate Cu(I) ions within *Mt*Csp1 are conserved in the *N. gonorrhoeae* protein. A model of the *N. gonorrhoeae* protein, based on the determined structure of *Mt*Csp1 [12], predicted it would adopt an identical fold (**Fig 2A**) in which all the 15 Cys thiols would be localized within the core of the four-helix bundle (**Fig 2B**). The model also predicted the protein would form a biological tetramer, like other Csp homologues [13]. Notably, the predicted tetramer structure resulted in localization of all predicted antibody epitopes on the surface of the complex exposed to solvent (**Supp Fig 1**), validating this structural model. Based on this sequence and structural homology, we hypothesized that NGO1701 encoded a Csp family protein.

**Figure 1.**
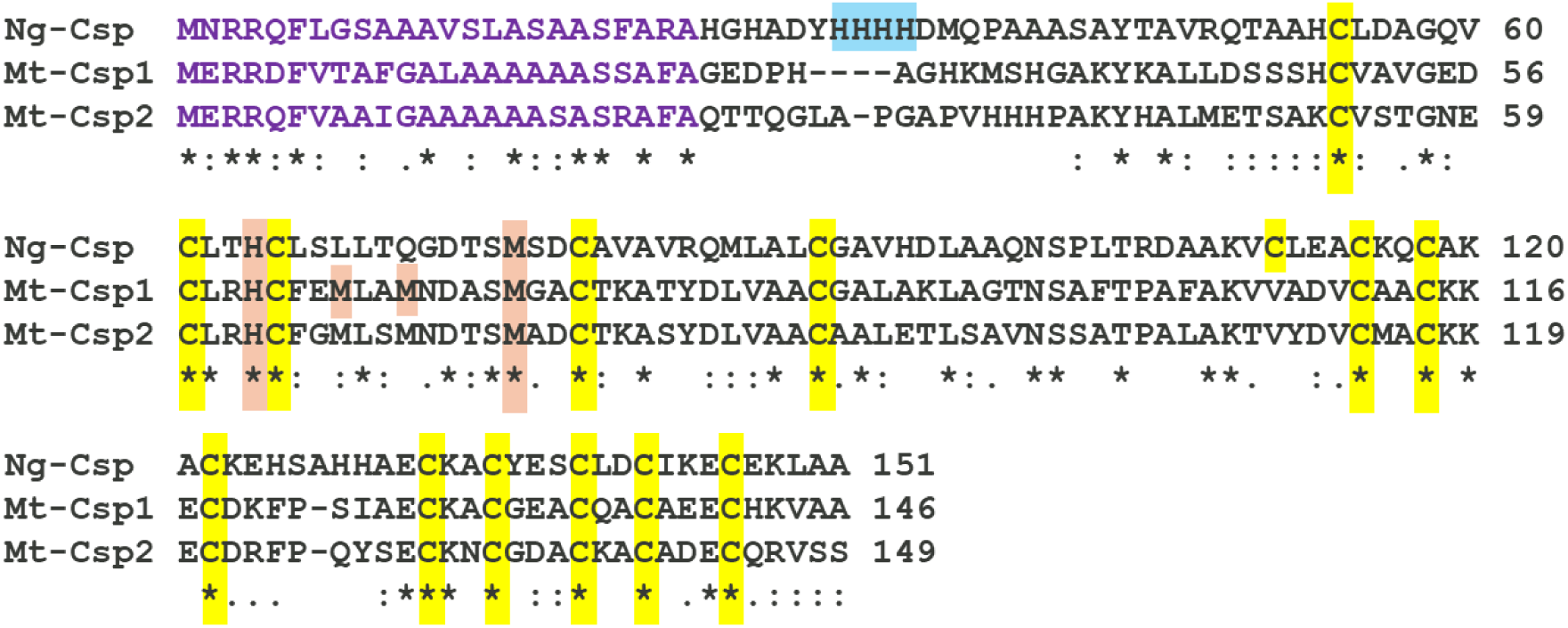
Sequence alignment of *N. gonorrhoeae* NGO1701 with related Csp proteins. Alignment of the NGO1701 protein sequence with two related sequences from *M. trichosporium* OB3b (*Mt*Csp1 and *Mt*Csp2) [12] with Clustal Omega. Putative twin arginine translocase (TAT) targeting pre-peptides are shown in violet, Cys residues highlighted in yellow, and Cu(I)-coordinating His and Met residues highlighted in orange. In NGO1701, additional N-terminal His residues potentially involved in copper coordination are shown in blue. Perfectly conserved residues are indicated by *, and highly or moderately conserved residues are indicated by the : and . symbols, respectively.

**Figure 2.**
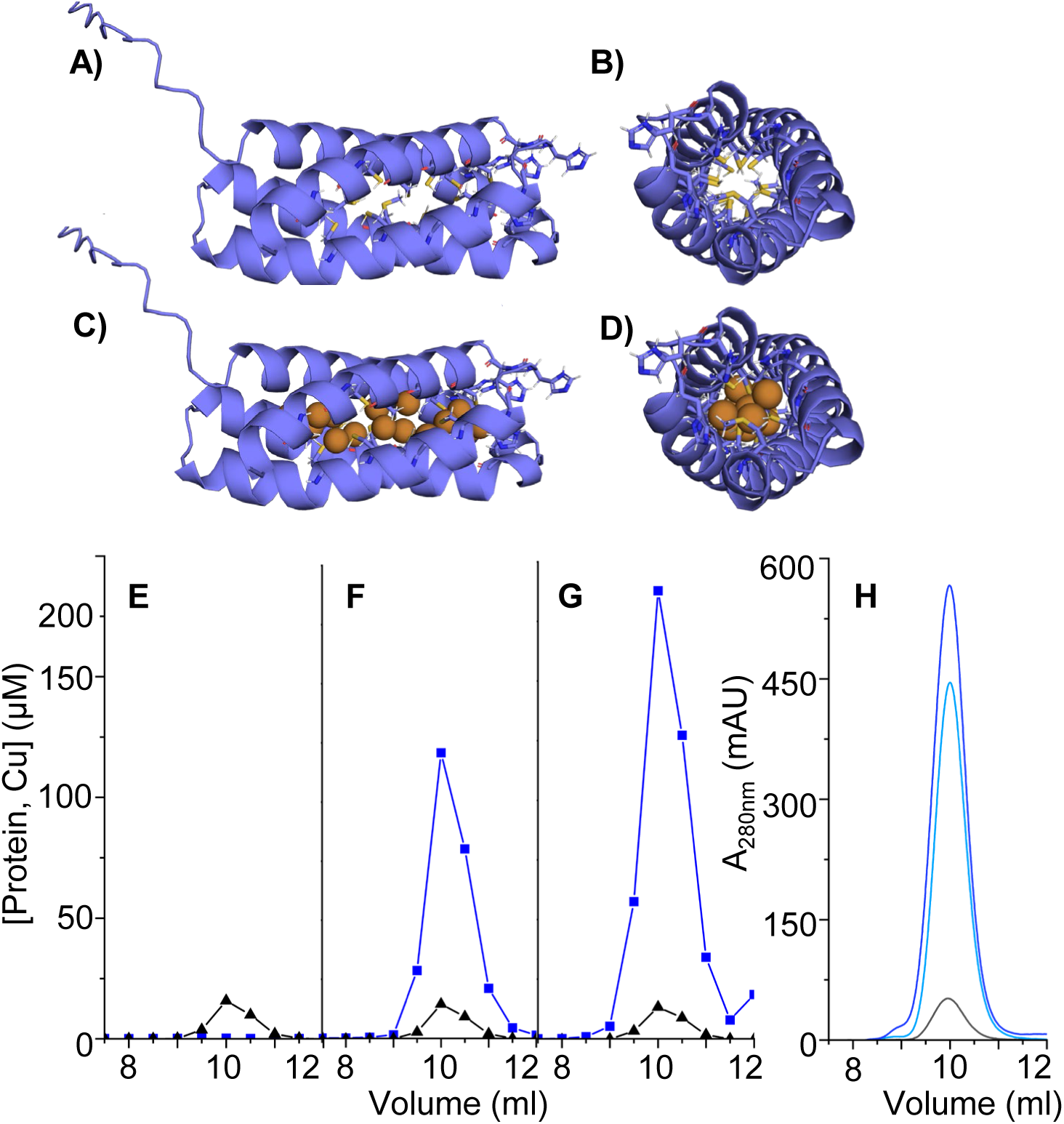
Structure model of Csp and copper binding. **(A-D)** Structural models of a *N. gonorrhoeae* Csp monomer produced using IntFOLD, based on the published structure of *Mt*Csp1 (PDB: 5FJD). **A)** Side-on, and **B)** end-on views of the apo-monomer, illustrating the predicted four-helix bundle structure (blue ribbon), with the metal-coordinating Cys, His and Met residues in ball-and-stick representation, colored by element (yellow, S; red, O; blue, N). **C)** Side-on, and **D)** end-on view of the same model loaded with 15 copper ions (bronze), using the same color scheme. **(E-H)** SEC analysis of purified recombinant apo-Csp **E)** before and after titration with **F)** 10 or **G)** 20 mole equivalents of Cu(I). Copper (blue) and protein concentration (black), calculated from S content, were determined by ICP-OES. **H)** Absorbance of the samples at 280 nm in **E-G)** eluted from the SEC column in the absence (black) or presence of 10 (light blue) or 20 (dark blue) mole equivalents of Cu(I) [12]

### The *N. gonorrhoeae* Csp protein binds copper ions with high stoichiometry

To determine whether the *N. gonorrhoeae* Csp homologue could function in an analogous manner to *Mt*Csp1, we first tested whether the protein shares its proposed function, i.e. the ability to bind large numbers of Cu(I) ions. Recombinant Csp purified from *E. coli* BL21 cultured in LB medium was devoid of bound copper (<0.1 mole equivalent detected) or any other metal ion by ICP-OES (**Fig 2E**). Incubation of recombinant Csp with 10 or 20 mole equivalents of Cu(I) *in vitro*, followed by separation of unbound Cu by SEC (**Fig 2F** and **2G, respectively**), demonstrated that Csp bound copper in high stoichiometry, saturating at approximately 15 equivalents of copper associated with the protein. A dramatic increase in the protein’s absorbance at 280 nm after incubation with Cu(I) (**Fig 2H**) was consistent with the Cu(I) ions being primarily coordinated by Cys residues within the Csp tube, as predicted (**Fig 2C-2D**) and as previously demonstrated for *Mt*Csp1 [12]. Thus, the *N. gonorrhoeae* Csp binds a large number of Cu(I) ions, consistent with its homology to *Mt*Csp1, its predicted structure, and with a proposed role in *N. gonorrhoeae* copper homeostasis.

### Allelic analysis and conservation of Csp in *Neisseria* sp

We previously showed that the gene encoding Csp is conserved in *N. gonorrhoeae* [8]. Since our original analysis, the number of gonococcal genomes available in the PubMLST database has increased to 25,553 [15]. A new analysis of the *csp* (*NEIS2720*) gene sequence confirmed conservation in 48 alleles identified, with allele 1 present in 20204 strains and allele 2 in 4135 strains (79% and 16% of all strains, respectively) (**Table 1**). Alleles that could not be assigned were present in 653 *N. gonorrhoeae* strains (∼2.5% of all strains), due to incomplete sequence data because the genes were located at the end of contigs; *csp* was present on allele 16 in 408 strains (1.6% of strains) and on several other alleles in 15 strains total (**not shown**). No polymorphic sites were detected, indicating protein sequence conservation (**Table 1**). Analysis of presence and conservation of *csp* was also carried out in other pathogenic and non-pathogenic *Neisseriae* genomes. The gene was present in *N. lactamica* (65 alleles in 1196 strains), *N. polysaccharea* (19 alleles in 86 strains), *N. bergeri* (6 alleles in 62 strains) and *N. cinerea* (9 alleles in 47 strains) (**Table 1**), with several polymorphic sites and some consistent non-synonymous amino acid sequence mutations among these species. Notably, none of these mutations altered the residues predicted to bind metal ions in Csp. Interestingly, no record of *csp* presence in *N. meningitidis* and *N. subflava* was found in PubMLST (**Table 1**), although a recent report has indicated its presence in the latter (WP_137041264.1) [16]. Thus, despite being present in multiple species, Csp has potential as a *N. gonorrhoeae*-specific vaccine target.

**Table 1.**
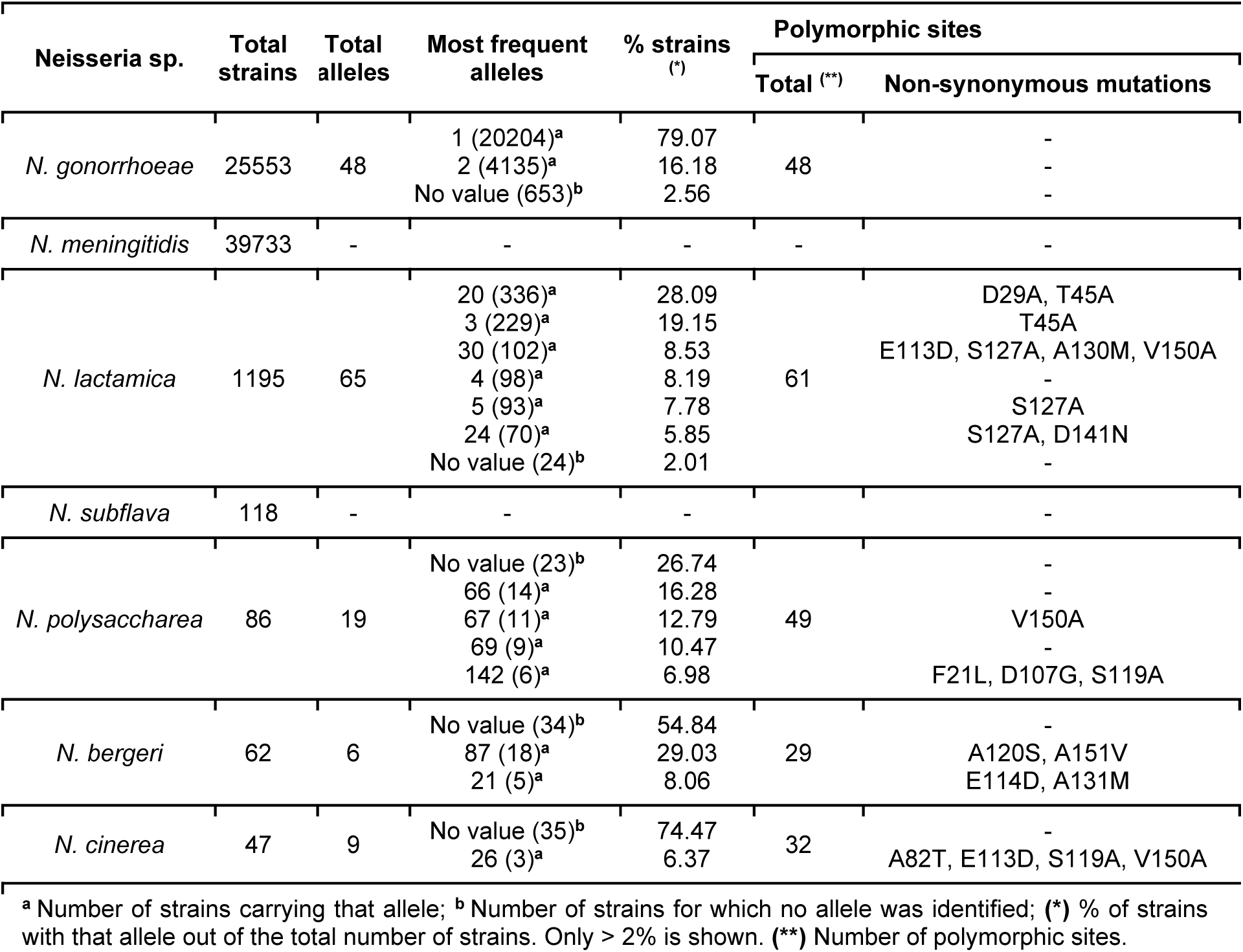
*csp* (NEIS2720) allelic profile analysis from PubMLST.

| Neisseria sp. | Total strains | Total alleles | Most frequent alleles | % strains (*) | Polymorphic sites |  |
| --- | --- | --- | --- | --- | --- | --- |
|  |  |  |  |  | Total (**) | Non-synonymous mutations |
| *N. gonorrhoeae* | 25553 | 48 | 1 (20204)<sup>a</sup> | 79.07 | 48 | - |
|  |  |  | 2 (4135)<sup>a</sup> | 16.18 |  | - |
|  |  |  | No value (653)<sup>b</sup> | 2.56 |  | - |
| *N. meningitidis* | 39733 | - | - | - | - | - |
| *N. lactamica* | 1195 | 65 | 20 (336)<sup>a</sup> | 28.09 | 61 | D29A, T45A |
|  |  |  | 3 (229)<sup>a</sup> | 19.15 |  | T45A |
|  |  |  | 30 (102)<sup>a</sup> | 8.53 |  | E113D, S127A, A130M, V150A |
|  |  |  | 4 (98)<sup>a</sup> | 8.19 |  | - |
|  |  |  | 5 (93)<sup>a</sup> | 7.78 |  | S127A |
|  |  |  | 24 (70)<sup>a</sup> | 5.85 |  | S127A, D141N |
|  |  |  | No value (24)<sup>b</sup> | 2.01 |  | - |
| *N. subflava* | 118 | - | - | - | - | - |
| *N. polysaccharea* | 86 | 19 | No value (23)<sup>b</sup> | 26.74 | 49 | - |
|  |  |  | 66 (14)<sup>a</sup> | 16.28 |  | - |
|  |  |  | 67 (11)<sup>a</sup> | 12.79 |  | V150A |
|  |  |  | 69 (9)<sup>a</sup> | 10.47 |  | - |
|  |  |  | 142 (6)<sup>a</sup> | 6.98 |  | F21L, D107G, S119A |
| *N. bergeri* | 62 | 6 | No value (34)<sup>b</sup> | 54.84 | 29 | - |
|  |  |  | 87 (18)<sup>a</sup> | 29.03 |  | A120S, A151V |
|  |  |  | 21 (5)<sup>a</sup> | 8.06 |  | E114D, A131M |
| *N. cinerea* | 47 | 9 | No value (35)<sup>b</sup> | 74.47 | 32 | - |
|  |  |  | 26 (3)<sup>a</sup> | 6.37 |  | A82T, E113D, S119A, V150A |

<sup>a</sup> Number of strains carrying that allele; <sup>b</sup> Number of strains for which no allele was identified; (\*) % of strains with that allele out of the total number of strains. Only > 2% is shown. (\*\*) Number of polymorphic sites.

### Construction and immune characterization of *N. gonorrhoeae csp* deletion mutant and complemented strains

To investigate the function of Csp in *N. gonorrhoeae*, a *ngo1701* deletion mutant strain was generated (Δ*csp*) in *N. gonorrhoeae* F62, and a corresponding *ngo1701* complemented strain (*csp_c).* Presence and expression of Csp was examined by dot blot using sera from mice previously immunized with purified NGO1701 [8]. *N. gonorrhoeae* F62, Δ*csp* and *csp_c* (uninduced) were grown on GC plates and in GCB medium without IPTG; to induce Csp expression, *csp_c* was grown on plates containing 100 µM IPTG but inoculated in broth with increasing concentrations of IPTG (0, 0.25 mM, 0.5 mM or 1 mM). After 2 h incubation, the cultures were diluted and equivalent numbers of cells (approx. 7.5x10^5^ bacteria) were spotted on nitrocellulose filters. Csp expression was detected in *csp_c,* both when uninduced (pGCC4 is a known leaky plasmid [17]) and on plates containing IPTG (**Fig 3A, dots 2 and 3**), and it was increased by addition of IPTG also in liquid culture (**Fig 3A, dots 4-6**). A final IPTG concentration of 0.25 mM was chosen for all subsequent experiments and bacteria grown in these conditions are referred to as *csp_c+*, while bacteria grown in liquid medium without IPTG are referred to as *csp_c.* A low-level immunoreactivity of Δ*csp* with anti-NGO1701 mouse sera was observed (**Fig 3A, dot 7**), and of all strains with sera from mice immunized with alum alone (**Fig 3B, dots 1-7**), which were consistent with non-specific sera reactivity with bacteria. Purified recombinant Csp (50 ng) was only recognized by the anti-NGO1701 mouse sera (**Fig 3A-B, dots 8**), indicating antigen specificity.

**Figure 3.**
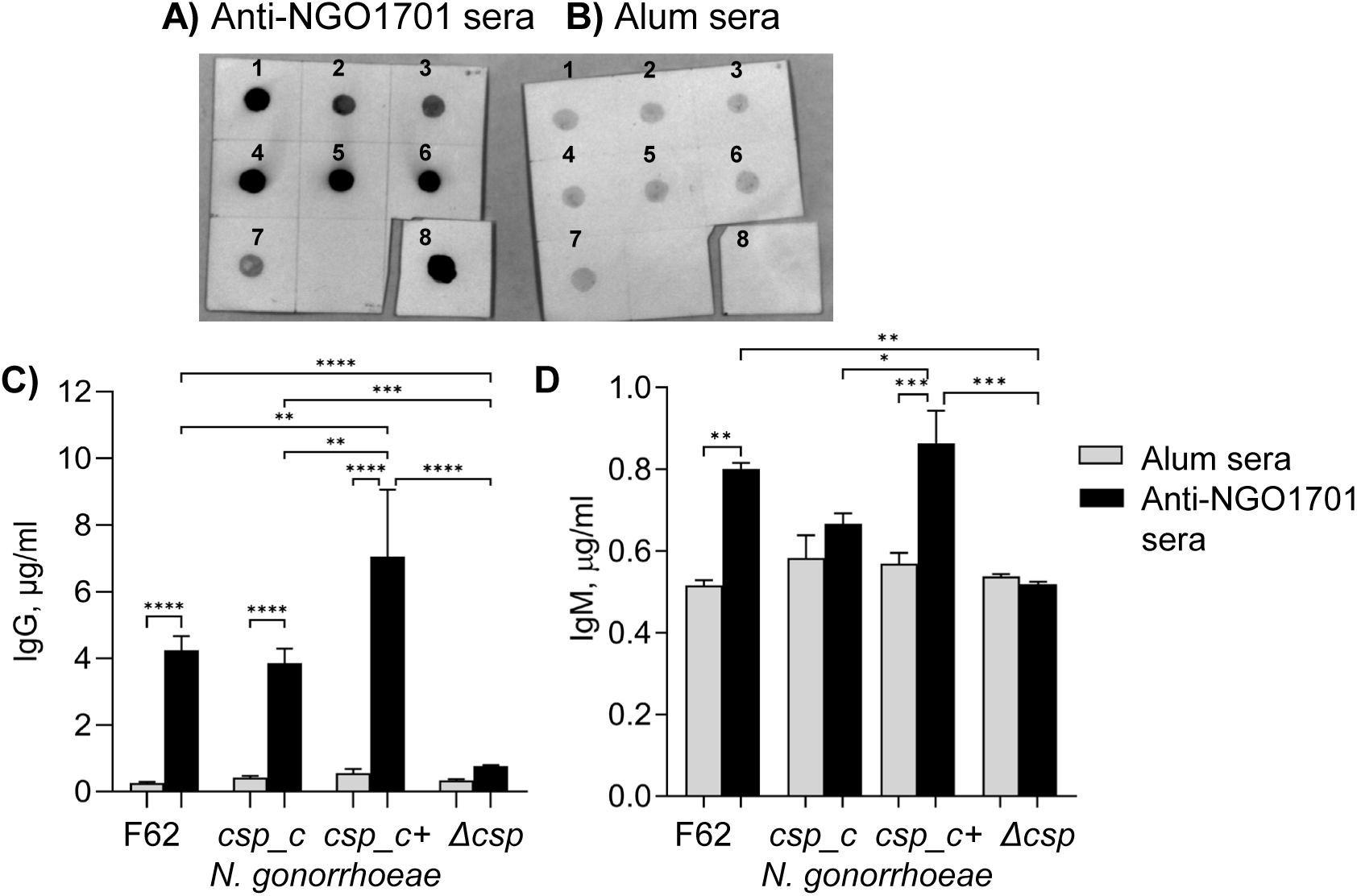
Csp protein expression in *N. gonorrhoeae* and immune recognition. Bacterial suspensions (OD_600_ 0.33) were spotted on nitrocellulose (5 μl/dot) and examined by dot blot with **A)** pooled sera from mice immunized with NGO1701 and alum or **B)** alum alone [8] (1:1000 dilution). **1)** *N. gonorrhoeae* F62 wildtype; **2)** uninduced *csp_c*; **3)** *csp_c* plated on GC agar plates containing 100 μM IPTG and grown in GCB without IPTG; **4)** *csp_c* plated as above and grown in GCB with 0.25 mM IPTG, **5)** 0.5 mM IPTG or **6)** 1 mM IPTG; **7)** Δ*csp;* **8)** purified recombinant Csp (50 ng). **C)** Total IgG antibodies (µg/ml ± SEM) measured by ELISA of mouse sera as above against *N. gonorrhoeae* F62 wildtype, Δ*csp, csp_c* and *csp_c+* (0.25 mM IPTG). Alum alone sera (gray bars), anti-NGO1701 and alum (black bars). Sera were tested in triplicate or quadruplicate. **D)** Total IgM antibodies as in **C)**. * p = 0.05, ** p = 0.005, *** p = 0.0001 and **** p < 0.0001 by one way ANOVA with Tukey’s comparison test.

A whole-cell ELISA was used to quantify antibody recognition of Csp. Similar levels of IgG antibodies recognizing *N. gonorrhoeae* F62 wildtype and *csp_c* were detected, confirming baseline Csp expression in both strains, and higher IgG antibody levels against *csp_c+* confirmed protein induction (**Fig 3C**). Consistent with the dot blot results, immune recognition of Δ*csp* was significantly lower **(Fig 3C**), comparable to non-specific immunoreactivity of sera from mice immunized with alum alone (**Fig 3C, gray bars**). Total IgM antibody levels were also measured, showing significantly higher reactivity against *N. gonorrhoeae* F62, *csp_c* and *csp_c+* than against Δ*csp* (**Fig 3D, black bars**), which was, again, comparable to IgM levels in sera from mice immunized with alum alone (**Fig 3D, gray bars**).

Phenotypic characterization of Csp in *N. gonorrhoeae in vitro*.

We investigated the role of Csp in gonococcal physiology by examining the growth kinetics of *N. gonorrhoeae* F62 wildtype, Δ*csp* and *csp* complemented cells. No difference among the strains was observed in colony size or morphology by phase-contrast light microscopy of plated bacteria (**Fig 4A-D**). Growth in liquid medium was monitored for 6 h; no major difference was observed in growth of Δ*csp* by optical density (**Fig 4E, squares**), except for a slightly lower OD_600_ at 5 h compared to the wildtype strain (**Fig 4E circles**) and at 6 h compared to *csp_c* (**Fig 4E, triangles**). In addition, no significant difference in the number of Δ*csp* colony forming units (CFU)/ml was observed compared to *N. gonorrhoeae* F62 wildtype (**Fig 4F, squares and circles, respectively**) or *csp_c* (**Fig 4F, triangles**) throughout the 6 h growth curve. Growth kinetics of *csp_c+* showed a significantly higher number of CFU/ml at 4, 5 and 6 h (**Fig 4F, diamonds**) but no significant difference in OD_600_ (**Fig 4E, diamonds**). These results suggested that Csp does not play a role in *N. gonorrhoeae* viability and fitness *in vitro*, but that IPTG induction of Csp improved bacterial growth.

**Figure 4.**
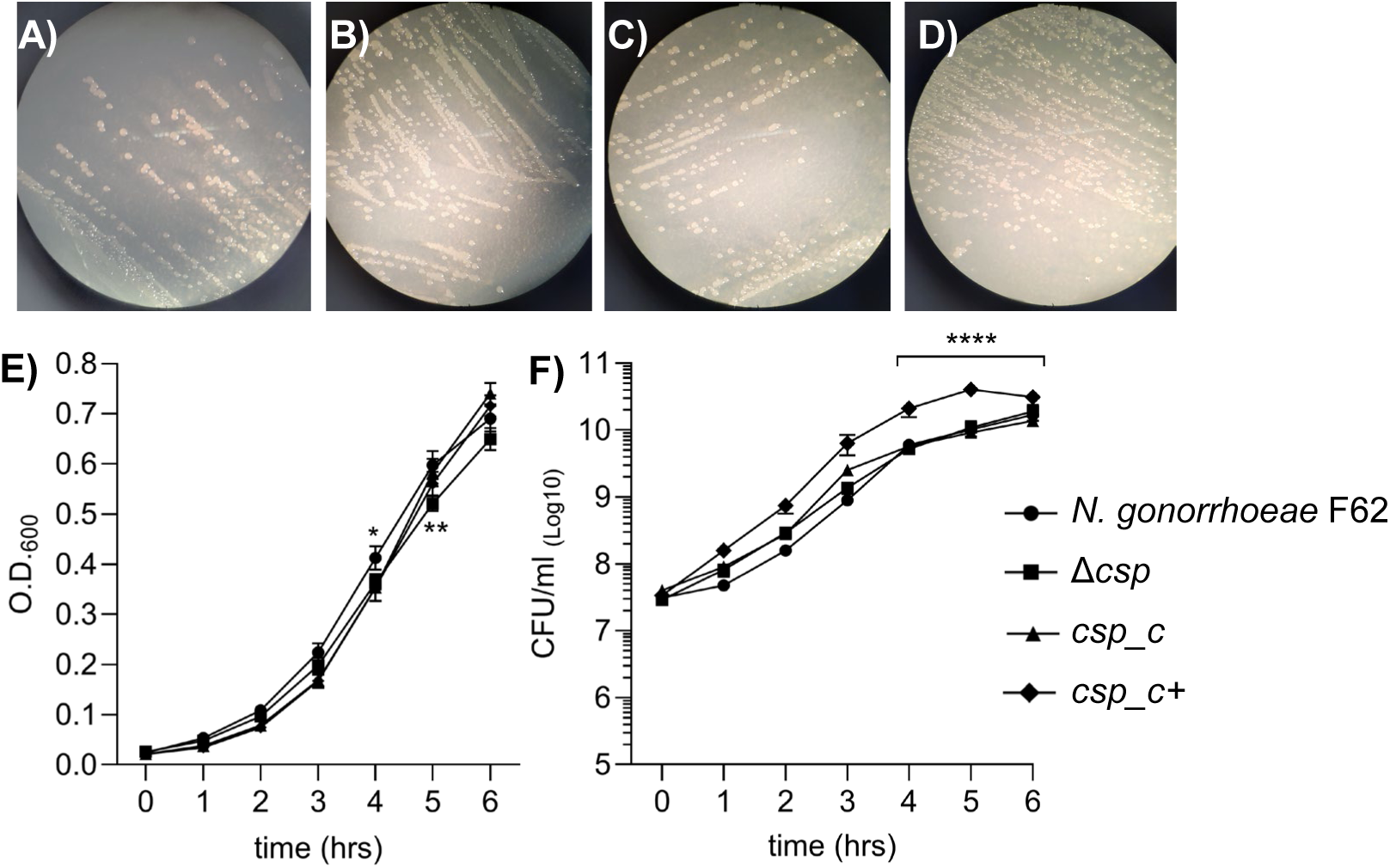
Colony morphology and growth curves. Light microscopy images showing colony morphology of **A)** *N. gonorrhoeae* F62 wildtype, **B)** Δ*csp*, **C)** *csp_c* and **D)** *csp_c+* on GC agar plates. **E)** OD_600_ values (average ± SEM) and **F)** Colony forming units (CFUs)/ml (average ± SEM) from >10 individual experiments per each strain. *N. gonorrhoeae* F62 wildtype (circles), Δ*csp* (squares), *csp_c* (triangles) and *csp_c+* (diamonds). 4 h, * p < 0.05 (*csp_c* and *csp_c+* vs wildtype); 5 h, ** p < 0.005 (Δ*csp* vs wildtype) and **** p < 0.0001 by 2way ANOVA with Dunnett’s multiple comparisons test.

### Copper-mediated Csp function in *N. gonorrhoeae*

A putative role of Csp in copper homeostasis in *N. gonorrhoeae* was examined by analyzing growth of the mutant bacteria in the presence of increasing concentrations of copper sulfate (CuSO_4_). In *N. gonorrhoeae* F62 wildtype, a decrease in OD was observed starting at 100 µM copper, compared to control cultures with no copper added (**Fig 5A, dark gray circles and black circles, respectively**). Significantly lower OD_600_ values were measured with 200 µM copper at the 4, 5 and 6 h time points (**Fig 5A light gray circles**), and growth stopped in the early exponential phase when bacteria were exposed to 500 µM copper (**Fig 5A, open circles**). This growth defect was also reflected in the number of CFU/ml of the bacterial suspensions in the presence of copper (**Fig 5E**). Copper-mediated toxicity was amplified by deletion of Csp. Growth of Δ*csp* in the presence of 200 µM and 500 µM copper (**Fig 5B** and **5F, light gray and open squares**) was significantly lower compared to Δ*csp* grown in the absence of copper (**Fig 5B** and **5F, black squares**) and to *N. gonorrhoeae* F62 wildtype at the same copper concentrations. Bacterial survival was rescued in *csp_c* to levels comparable to the wildtype strain (**Fig 5C** and **5G**) and further improved for *csp_c+* (**Fig 5D** and **5H**).

**Figure 5.**
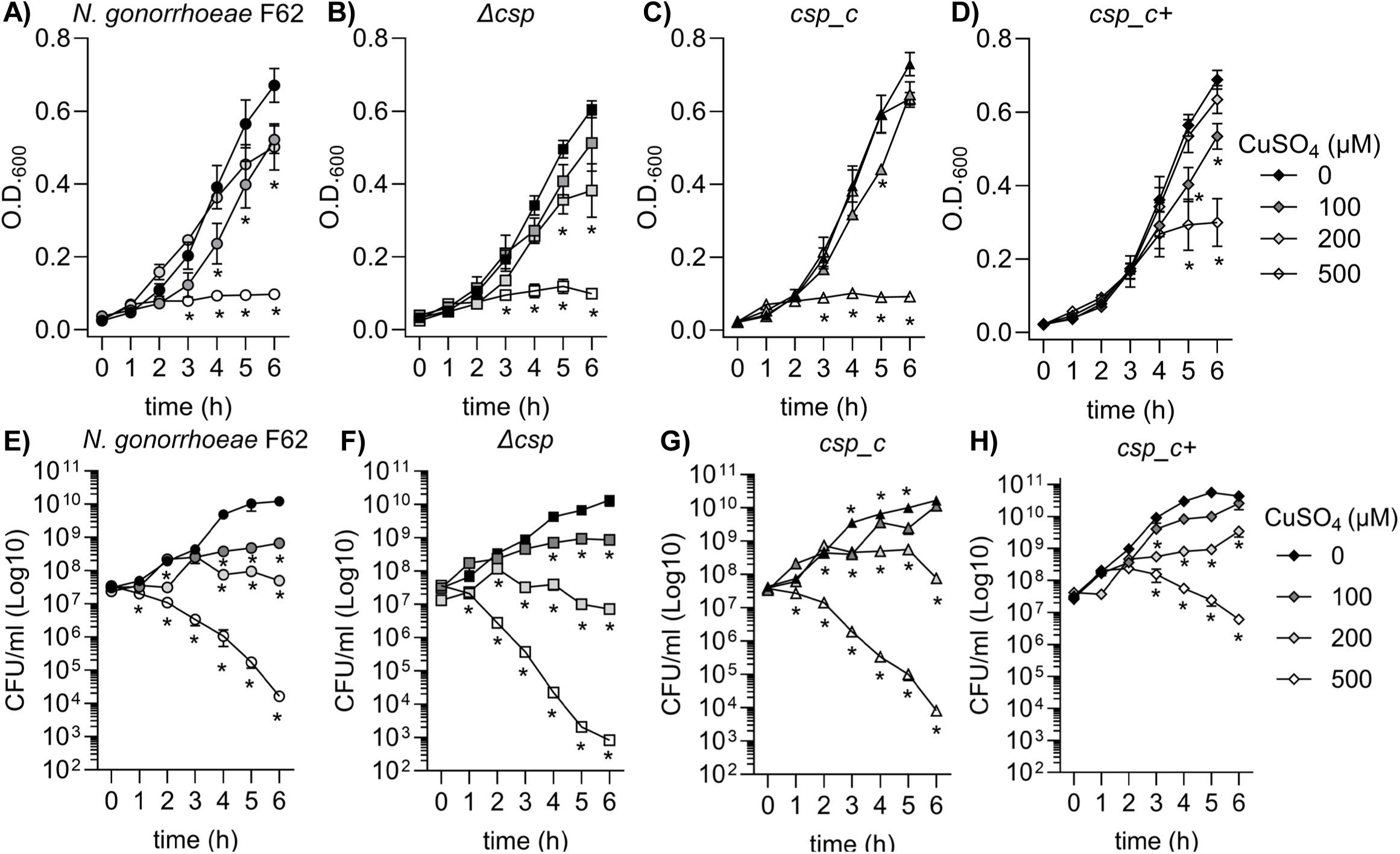
Dose-dependent effect of copper on bacterial growth. *N. gonorrhoeae* F62 wildtype (circles), Δ*csp* (squares), *csp_c* (triangles) and *csp_c+* (diamonds) were grown for 6 h in the presence of 100 µM copper sulfate (CuSO_4_) (dark gray symbols), 200 µM (light gray symbols), 500 µM (open symbols) or without copper (black symbols). **A-D**) OD_600_ values (average ± SEM) from a minimum of three independent experiments for each strain. * p ≤ 0.05 vs no copper by 2-way ANOVA with Dunnett’s multiple comparisons test. **E-H)** CFUs/ml (average ± SEM) as above. Statistical significance was determined by multiple Mann-Whitney test with Holm-Sidak correction set for p = 0.05 vs no copper for each strain, indicated by *.

Additional evidence for the role of Csp in *N. gonorrhoeae* resistance to copper was obtained using a disc diffusion assay. Bacterial suspensions were plated as a lawn on GC agar plates and exposed to paper discs impregnated with different concentrations of sterile CuSO_4_ solution. A dose-dependent zone of inhibition around the discs was visible for *N. gonorrhoeae* F62 wildtype. The highest copper concentration (200 mM) (**Fig 6A, top**) led to a wider area devoid of bacteria, which became smaller as the copper concentration decreased (**Fig 6A**). No zone of inhibition was visible around the control disc soaked with water (**Fig 6A, center**). Δ*csp* showed a wider zone of inhibition at all copper concentrations (**Fig 6B**), confirming increased sensitivity to copper toxicity. Resistance to copper of both *csp_c* and *csp_c+* was restored to levels similar to the wildtype strain (**Fig 6C** and **6D, respectively**). The size of the zone of inhibition for each strain was quantified (**Fig 6E**), confirming higher sensitivity of Δ*csp* to copper toxicity. Collectively, these results supported a role for Csp in copper resistance in *N. gonorrhoeae*.

**Figure 6.**
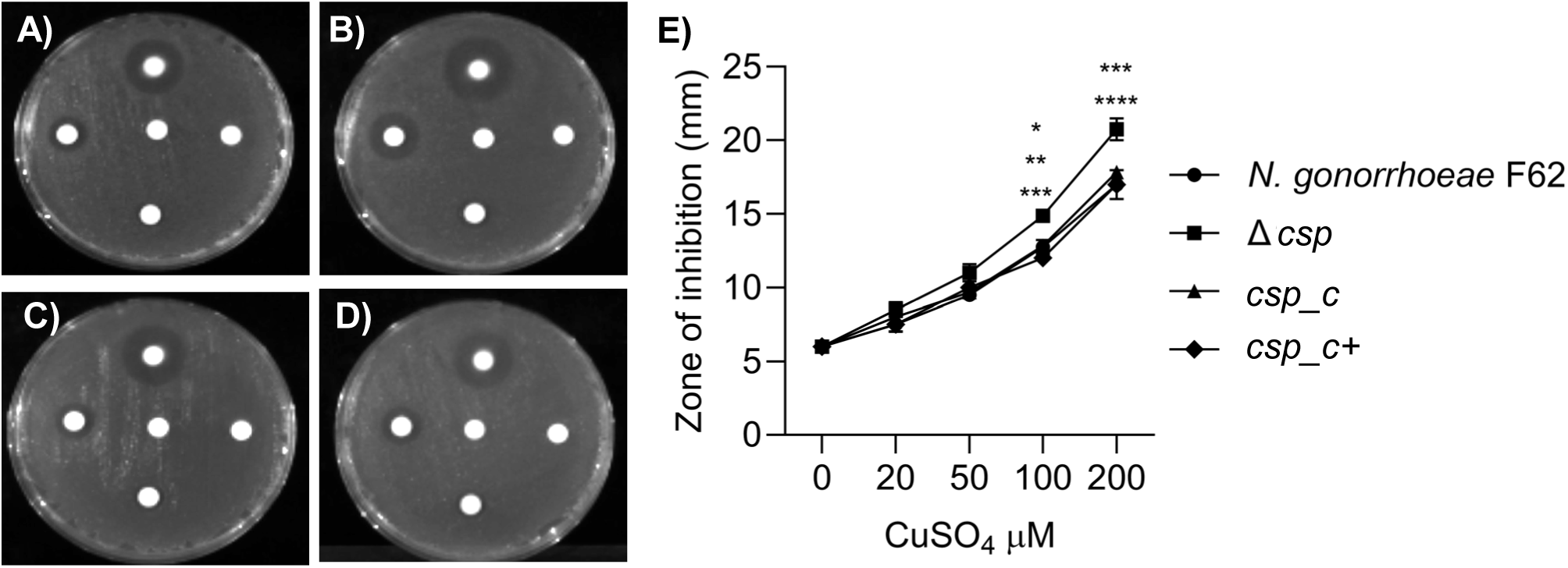
Disc diffusion assay. Representative images of the zone of inhibition around discs presoaked with increasing concentrations of CuSO_4_ (counterclockwise from top: 200 mM, 100 mM, 50 mM and 20 mM) or sterile water (center). **A)** *N. gonorrhoeae* F62 wildtype, **B)** Δ*csp,* **C)** *csp_c* and **D)** *csp_c+*. **E)** The diameter of the zone of inhibition was measured in mm and expressed as average ± SD from triplicate experiments. *N. gonorrhoeae* F62 wildtype (circles), Δ*csp* (squares), *csp_c* (triangles) and *csp_c+* (diamonds). * p = 0.013, ** p < 0.005, *** p < 0.0005 and **** p < 0.0001 by 2way ANOVA with Dunnett’s multiple comparisons test.

Other transition metals important for bacterial growth and functions were examined. No difference in sensitivity to nickel (NiSO_4_, 200 μM) between *N. gonorrhoeae* F62 and Δ*csp* was observed, with similar OD_600_ values and CFU numbers throughout the 6 h growth curves (**Supplemental Fig 2A-B)**. This suggested no role for Csp in resistance to nickel toxicity. Similarly, no differences in bacterial growth were observed when Δ*csp* was exposed to either 100 µM ferric nitrate (iron-replete) or 100 μM desferal (iron-deplete) compared to *N. gonorrhoeae* F62 wildtype (**not shown**), nor to excess zinc (50 μM ZnCl_2_) (**not shown**). Sensitivity to manganese (25 µM MnSO_4_) (**Supplemental Fig 2C-D**) and cobalt (50 µM CoCl_2_) (**Supplemental Fig 2E-F**) appeared to be slightly increased by Csp deletion, with a statistically significant lower OD_600_ and number of CFUs than wildtype *N. gonorrhoeae* F62 starting at the 5 h or 3 h time point, respectively. Resistance to cobalt was restored in *csp_c* and *csp_c+* to (**Supplemental Fig 2E-F**), suggesting that Csp might also be involved in buffering excess of these metals. However, incubation of recombinant Csp with even 1 mole equivalent of Co(II) led to protein precipitation (**not shown**), suggesting that these ions are unlikely to be able to enter the Csp tube. We conclude, therefore, that these other metal phenotypes are likely to be indirect, independent of direct buffering of these metal ions by Csp inside *N. gonorrhoeae* cells. Together, these results suggested that Csp was not likely to be involved in buffering of metal ions other than Cu(I), supporting a copper-specific function for Csp in *N. gonorrhoeae*.

### Intracellular copper measurement

Copper accumulation by *N. gonorrhoeae* F62 wildtype and Δ*csp* was also assessed. Bacteria were digested, and the elemental composition of the resulting lysate was determined by ICP-OES. After adjusting the metal content for sulfur content to normalize for biomass, the results showed that Δ*csp* contained approximately 60% of the cellular copper content of *N. gonorrhoeae* F62 wildtype after 4 h culture under basal copper conditions (**Fig 7**). No significant decreases were detected in cellular content of zinc (**Fig 7**), manganese, iron or magnesium (**Supplemental Fig 3**). No difference in cellular copper content was detected between *N. gonorrhoeae* F62 wildtype and Δ*csp* cells cultured for 4 h in the presence of 200 μM copper (**Fig 7**). These data support a model in which Csp stores copper in *N. gonorrhoeae*, presumably for supply of this cofactor to copper-dependent enzymes in the cell envelope.

**Figure 7.**
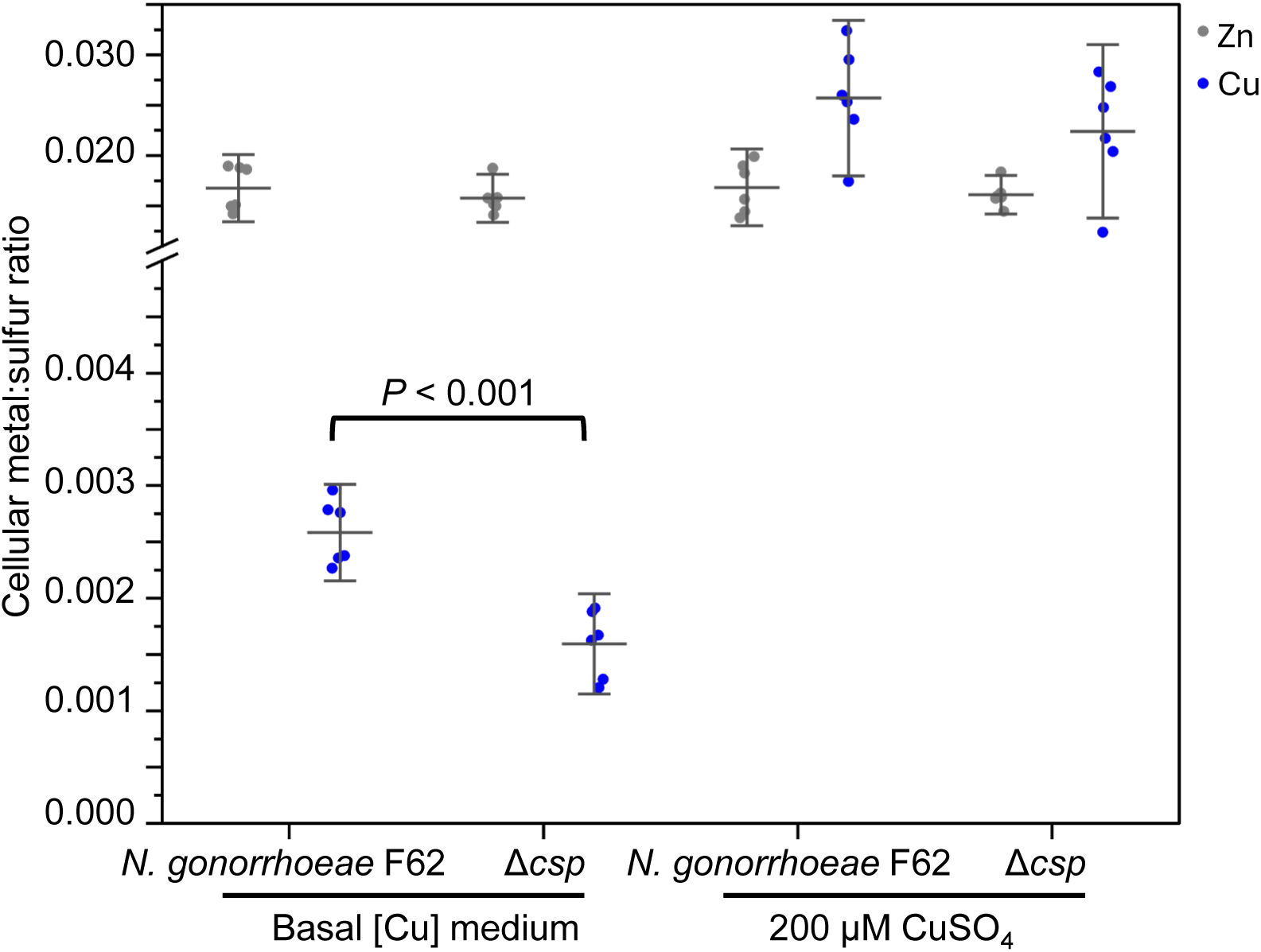
Copper accumulation is lower in *N. gonorrhoeae* Δ*csp* under basal conditions. *N. gonorrhoeae* F62 wildtype and Δ*csp* were grown for 4 h in GCB without added copper (basal [Cu] medium) or in the presence of 200 μM copper sulphate. After removing surface-adsorbed metal, bacterial pellets were digested and examined for Cu (blue), Zn (grey) and S content by ICP-OES. Metal content was normalized to S content to account for differences in biomass. *P* represents the results of an unpaired *t* test.

### Csp does not play a direct role in *N. gonorrhoeae* interaction with host epithelial cells

We evaluated whether presence and expression of Csp impacted the ability of *N. gonorrhoeae* to adhere to and invade host cells using HeLa cells, a human epithelial reproductive tract cell line previously shown to support gonococcal invasion [18–21]. HeLa cells were incubated with live *N. gonorrhoeae* F62, Δ*csp*, *csp_c* or *csp_c+* at a multiplicity of infection (MOI) of 100 for 2 h; monolayers were washed, lysed and plated for colony counting to quantify the number of cell-associated bacteria. No difference was detected among the strains (**not shown**). To evaluate internalized bacteria, a gentamicin protection assay was used. HeLa cells were incubated with *N. gonorrhoeae* as above, unattached bacteria were removed, and medium containing gentamicin was added to kill adherent bacteria. After overnight incubation without antibiotic overnight, the total number of intracellular bacteria was also quantified by colony counting, and no difference was observed among the strains (**not shown**). These results suggested that Csp did not play a direct role in gonococcal adhesion/invasion processes. Next, to evaluate whether Csp affects host cell activation, production of the pro-inflammatory cytokine IL-8 was used as read-out. HeLa cells (10^4^/ml) were incubated with *N. gonorrhoeae* F62, Δ*csp*, *csp_c* or *csp_c+* strains at MOIs of 10 and 100 overnight, and IL-8 secretion was measured by ELISA of the co-culture supernatants. To prevent bacterial proliferation during the incubation, formalin-inactivated *N. gonorrhoeae* were used. All strains induced similar levels of IL-8 secretion in a dose-dependent manner according to the bacterial MOI (**Supplemental Fig 4A** and **4B**). Although a small, statistically significant lower IL-8 production was observed in response to Δ*csp* compared to *N. gonorrhoeae* F62 wildtype, *csp_c* and *csp_c+*, this was unlikely to be biologically significant, suggesting that presence and expression of Csp had a minimal impact on IL-8 induction. Purified Csp (10 µg/ml) also induced low levels of IL-8 secretion (**Supplemental Fig 4A** and **4B, gray bars**). Overall, these results did not suggest a role for Csp in *N. gonorrhoeae* interaction with host cells *in vitro*.

### Serum bactericidal activity (SBA)

We previously showed that anti-NGO1701 mouse sera have bactericidal activity against several *N. gonorrhoeae* strains, with killing titers ranging from 1/5 to 1/40, depending on the strain [8]. When anti-NGO1701 mouse sera were tested against *N. gonorrhoeae* Δ*csp*, killing was almost completely abrogated compared to *N. gonorrhoeae* F62 wildtype (**Fig 8A** and **8B**), even at the highest sera concentration used (1/10). SBA was restored against *csp_c* (**Fig 8C**) and *csp_c+* (**Fig 8D**). These results supported our previous studies and the potential of Csp as a vaccine candidate antigen against *N. gonorrhoeae*.

**Figure 8.**
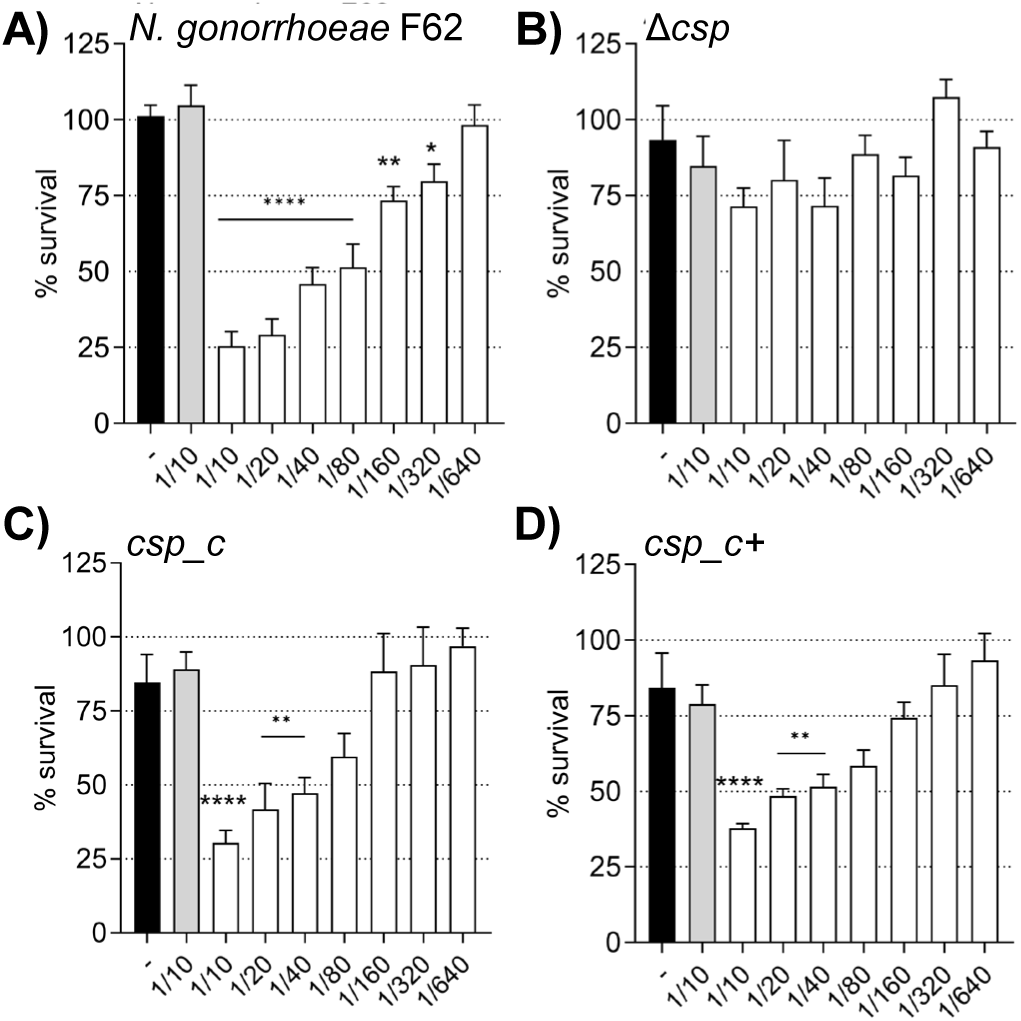
Serum bactericidal activity (SBA). Survival of **A)** *N. gonorrhoeae* F62 wildtype, **B)** Δ*csp,* **C)** *csp_c* and **D)** *csp_c+* incubated with sera from mice immunized with alum alone (gray bars) or NGO1701 and alum (white bars), expressed as % CFU (T30/T0) ± SEM from triplicate experiments. Bacteria alone, (black bars). * p < 0.05, ** p < 0.009 and **** p < 0.0001 by ordinary one-way ANOVA with Dunnett’s multiple comparisons test vs adjuvant alone sera. Sera dilutions are shown on the x-axis.

## Discussion

Rising antimicrobial resistance in *N. gonorrhoeae* has sparked interest in developing a vaccine against this sexually transmitted infection to mitigate the global disease burden. Initial clinical trial studies with a killed whole-cell vaccine or purified antigens (pilin, porins) failed to confer protection against heterologous re-infection in a human volunteer male urethral infection model. This was attributed to antigenic variability and induction of blocking antibodies (reviewed in [5]). Recently, outer membrane vesicle (OMV)-based vaccines developed for *N. meningitidis* have suggested cross-protective potential against *N. gonorrhoeae* [22, 23], supported by induction of bactericidal antibodies and protection in preclinical studies [24–26]. Gonococcal OMVs have also been tested, showing serogroup-specific protection in a mouse model of gonococcal vaginal colonization [27]. However, the protein content of the OMVs is just beginning to be characterized and it is still unclear how many and which targets are responsible for the observed protection [28, 29].

NGO1701 is a gonococcal hypothetical protein that we identified as a potential vaccine antigen through a novel transcriptomics-based approach (CASS). Immunization of mice with this protein (alone, or combined with two other CASS candidates, NGO0690 and NGO0948 (BamC)) induced antibodies with bactericidal activity [8, 11]; serum bactericidal activity (SBA) is regarded as the closest *in vitro* surrogate of protection assay for *N. gonorrhoeae* [30]. In ongoing experiments, immunization with the trivalent vaccine and Alum+MPLA as adjuvant has shown a significant reduction in bacterial burden and a faster gonococcal clearance in a mouse model of gonococcal vaginal colonization (*P. Massari and S. Ram, unpublished*). A bivalent vaccine (NGO1701+NGO0690) and even NGO1701 alone were also protective, albeit to a lower extent than the three-antigen vaccine (*P. Massari and S. Ram, unpublished*). However, due to lack of information about NGO1701 function in *N. gonorrhoeae*, it is impossible to predict whether expression of this protein might be modified by *N. gonorrhoeae* as a mechanism of immune escape, while maintaining bacterial fitness *in vitro* and *in vivo*.

We have initiated the characterization of NGO1701, which we here designated as the *N. gonorrhoeae* copper storage protein (Csp). Consistent with sequence homology to characterized Csp proteins, we found that purified recombinant *N. gonorrhoeae* Csp was able to sequester approximately 15 Cu(I) ions per protein monomer. Copper was likely coordinated to Cys residues localized to the interior cavity of the putative four-helix bundle structure. Like a previously characterized Csp1 in *M. trichosporium* OB3b, the gonococcal Csp possesses a putative TAT signal sequence, consistent with its presence in the cell envelope but how it localizes to the bacterial surface, enabling recognition by antibodies, is unclear. A cellular function for Csp proteins in sequestration of excess copper to detoxify this highly redox active metal to prevent deleterious side-reactions has been tested in some systems but has yielded contradictory results. Heterologous expression of a cytosolic Csp in *E. coli* slightly increased growth [13], whereas a strain of *Streptomyces lividans* lacking its cytosolic Csp showed slightly reduced growth under toxic copper conditions [31]. However, a mutant strain of *B. subtilis* lacking its cytosolic Csp did not show inhibited growth in elevated copper [32]. Here, we have established that *N. gonorrhoeae* growth in high copper conditions correlates with Csp expression, with a Δ*csp* strain exhibiting growth inhibition and a *csp_c*+ strain with induced expression of Csp growing better relative to wildtype cells. Interestingly, we also observed reduced growth of Δ*csp* in the presence of other metals such as cobalt and manganese but, unlike Cu(I), we observed no sequestration of Co(II) or Mn(II) ions by the recombinant Csp protein. It seems likely that these phenotypes are caused by displacement of copper ions by excess cobalt or manganese within the cell envelope. Although the observed differences in growth under high copper conditions were small, they are notable given that Csp is secreted out of the cytosol. This indicates that sequestration of excess Cu(I) ions by Csp can protect some vulnerable target(s) in the cell envelope under copper stress conditions. This may be important during infection as copper detoxification is essential for pathogens to overcome copper toxicity during infection [14, 33, 34].

On the other hand, prior studies have indicated a homeostatic role for Csp proteins in the storage of copper cofactors for supply to copper-dependent enzymes. In *M. trichosporium*, Csp storage influences copper availability to the copper-sensing regulator that controls the switch between its copper-dependent and iron-dependent isozymes of methane monooxygenase [12], which performs the critical reaction in its utilization of methane. The cytosolic Csp in *Bacillus subtilis* has been suggested to supply copper to the multicopper oxidase enzyme CotA during sporulation [32]. *N. gonorrhoeae* lacking Csp accumulated significantly fewer copper atoms when cultured under basal copper conditions, suggesting a function for Csp in normal growth. *N. gonorrhoeae* possesses several important copper-dependent enzymes in the cell envelope, including cytochrome *cbb*^3^ oxidase, an azurin homologue, and nitric oxide reductase [35, 36]. The latter is both periplasmic and surface-localized, plays a role in virulence, and its metalation involves the cuprochaperone AccA [37]. Therefore, a working hypothesis is that Csp in the *N. gonorrhoeae* envelope can supply copper to one or more of these target copper-enzymes. Metals are important for gonococcal responses to host nutritional immunity and virulence but, since no considerable growth defect (i.e. colony morphology, size, optical density, CFU numbers and replication time) was observed for Δ*csp* compared to wildtype *N. gonorrhoeae* in standard culture conditions, it is possible that Csp is not critical for bacterial fitness. Whether Csp plays a role in *N. gonorrhoeae* virulence awaits explicit experimental testing. So far, we have not observed an effect of Csp on the ability of *N. gonorrhoeae* to adhere to and invade a human cervical epithelial cell line model, HeLa cells, widely used to study bacterial host cell interactions. While this result was not unexpected (there is no indication that Csp may act as an adhesin), future studies may examine other reproductive tract epithelial cell lines and immune cells, particularly macrophages. Similarly, no effect of Csp presence or expression by *N. gonorrhoeae* was observed on epithelial cell inflammatory responses *in vitro.* Unfortunately, analysis of the effect of copper on *N. gonorrhoeae* virulence in a host cell co-culture model is limited by the toxicity of copper in these systems; we observed a dose-dependent and time-dependent cytotoxic effect of copper on HeLa cells starting at 100 µM, a concentration for which only a small difference in survival was reported between Δ*csp* and *N. gonorrhoeae* F62 wild type. However, *csp* is indeed expressed *in vivo* by gonococci recovered from men and women naturally infected with *N. gonorrhoeae* [10, 38, 39], which was one of the attributes for its identification as a potential vaccine target. The ability of *N. gonorrhoeae* Δ*csp* to survive *in vivo* is currently being tested in a mouse model of vaginal colonization to correlate bacterial fitness with infection.

The bactericidal activity of anti-NGO1701 mouse sera is abrogated against Δ*csp*, and thus, if Csp expression by *N. gonorrhoeae in vivo* is turned down or suppressed as a vaccine escape strategy, the immune responses induced by vaccination with this antigen alone may become less relevant. However, including Csp in a multivalent subunit vaccine may contribute to broader protection and decrease the chances of vaccine failure. Understanding the precise role of Csp in gonococcal physiology and homeostasis, and especially its putative function in supply of copper to envelope copper-enzymes essential for virulence will enable better assessment of its future value as a therapeutic and vaccine target

## Materials and Methods

### Csp sequence analysis and structure modeling

The sequence of characterized *M. trichosporium* OB3b (*Mt*Csp1 and *Mt*Csp2) [12] Csp proteins were downloaded from NCBI and aligned with *N. gonorrhoeae* NGO1701 (new locus tag NGO_RS08465) using Clustal Omega [40] https://europepmc.org/article/MED/38597606. Annotations were added manually based on prior characterization studies [12, 13]. Structural modelling used IntFOLD and GalaxyHomomer [41, 42] and were visualized using Pymol [43] (available at: http://www.pymol.org/pymol). Allelic analysis of Csp (NEIS2720) in all *Neisseria sp*. Genomes available in the PubMLST database [15] was carried out to evaluate *csp* gene sequence presence and conservation.

### Expression of recombinant Csp in *E. coli*

*N. gonorrhoeae* Csp (Accession number: WP_003689877.1) was cloned into a pET plasmid, expressed in *E. coli* BL21 (DE3) cells in selective (50 μg/mL ampicillin) LB medium through 4 h induction with 1 mM IPTG, and initially purified by nickel affinity chromatography as previously described [8]. The purified protein was resolved by size exclusion chromatography (SEC) on a Superdex 75 10/300 GL column (Cytiva) in 20 mM Tris, pH 7.5, 150 mM NaCl, 5 mM EDTA. Purified protein was resolved by reducing SDS-PAGE on 16% (w/v) acrylamide gels and Coomassie staining confirmed the presence of a single band (Thermo).

### Construction of *csp* deletion mutant and complemented strains in *N. gonorrhoeae* F62

*N. gonorrhoeae* F62 wildtype (Pil+/Opa+) organisms were propagated from frozen glycerol stocks on GC medium base agar plates (Difco) supplemented with 1% (v/v) IsoVitaleX (GC plates) at 37°C with 5% CO_2_ overnight. Colonies were swabbed with a sterile loop and inoculated in liquid cultures in GC broth (Difco) supplemented with 1% IsoVitaleX (GCB) and grown at 37°C with shaking at 120 rpm for 2 h. Growth was measured spectrophotometrically at OD_600_ (OD_600_ = 1 corresponding to ∼ 1-2 x 10^9^ bacteria/ml). For generating a *csp* deletion mutant strain (Δ*csp*), the *ngo1701* gene was amplified by PCR using primers designed based on the available sequence of *ngo1701* from FA1090 (NCBI accession # NC_002946) (**Supplemental Table 1**). The chromosomal DNA region 428-bp upstream of *ngo1701* was amplified using upstream_F and upstream_R primers, and the region 262-bp downstream of *ngo1701* using downstream_F and downstream_R primers. Primers encoded *Eco*RI and *Hind*III sites, respectively, allowing the insertion of a kanamycin (Kan) resistance cassette (*kan*) containing the same restriction endonucleases. The Kan cassette (975-bp) was amplified from the pGCC4 plasmid (Addgene, Watertown, MA) using *kan*_F and *kan*_R primers, digested with *Eco*RI and HindIII (New England Biolabs, Ipswich, MA) and ligated with *ngo1701* upstream and downstream products single-digested with the same restriction enzymes. The resulting linear construct, *ngo1701::kan*, was transformed into naturally competent *N. gonorrhoeae* F62 and integrated into the chromosome by homologous recombination [44] to generate the Δ*csp* strain. Transformants were selected on GC plates containing Kan (50 μg/ml) and grown as described above. Correct insertion of *ngo1701::kan* into the chromosome was verified by PCR using a *ngo1701* upstream confirmation primer (*ngo1701*_F-check) and a primer (Kan_F/R) complementary to the Kan cassette (pGCC4 mid Kan), and by Sanger sequencing. All primers were synthesized by Integrated DNA Technologies (IDT) (Coralville, IA). For construction of the *ngo1701* complemented strain (*csp_c)*, pGCC4 plasmid construction was outsourced to Genscript (Piscataway, NJ). Complementation analysis employed the Neisseria insertional complementation system (NICS) [45]. All plasmids were confirmed by whole plasmid sequencing (Plasmidsaurus Inc.). Linearized pGCC4::*ngo1701* complementation plasmid was transformed into *N. gonorrhoeae* F62 wildtype and transformants selected on GC plates containing erythromycin (1.25 µg/ml). Genomic DNA from *N. gonorrhoeae* F62 harboring *ngo1701* at the complementation site (AspC-LctP) was used to transform *Δcsp*. Transformant colonies were selected on GC plates containing erythromycin (1.25 µg/ml) and kanamycin (100 µg/ml). The presence and proper insertion of the complementing copy of *ngo1701* between *lctP* and *aspC* was confirmed by PCR using LctP_F and Asp*C*_R primers or LctP_F and Laclq_R primers. PCR was also used to confirm mutation of the native chromosomal copy of *ngo1701*. The nucleotide sequence of the complementing copy of *ngo1701* in *N. gonorrhoeae csp_c* was confirmed by sequence analysis of the amplified PCR product (Genewiz, Azenta Life Sciences).

### Expression of Csp in *N. gonorrhoeae csp_c* and dot blot

The *csp_c* strain was grown on GC plates containing 100 µM IPTG. Bacteria were swabbed from the plates and inoculated in GCB containing 0.1, 0.25, 0.5 or 1 mM IPTG, or no IPTG. The *N. gonorrhoeae* F62 and the Δ*csp* strains were grown in GCB as previously described. Liquid cultures were grown for 2 h, diluted to OD_600_ = 0.33 and 5 μl aliquots (∼ 1.6 - 3.3 x 10^6^ bacteria total) were directly spotted onto nitrocellulose membranes. Purified recombinant Csp (5 µl, 50 ng total) was used as a positive control. Membranes were blocked with PBS/5% BSA and incubated overnight at 4°C with sera from mice immunized with alum alone or with NGO1701 and alum [8] (1:1000 dilution). Immunoreactivity was detected with an anti-mouse IgG secondary AP-conjugated antibody (Southern Biotech, Birmingham, AL, USA) and NBT/BCIP (5-bromo-4-chloro-3-indolyl phosphate/Nitroblue Tetrazolium) chromogenic substrate (Bio-Rad, Hercules, CA, USA). A concentration of 0.25mM IPTG in liquid culture was chosen for all subsequent experiments and bacteria grown in this condition were designated as *csp_c+*.

### Elemental analysis by inductively coupled plasma optical emission spectrometry (ICP-OES)

Recombinant Csp samples were digested in 500 μL concentrated (65% v/v) nitric acid (Merck) overnight at room temperature. Heat-killed bacterial samples were digested in 500 μL concentrated nitric acid at 70°C for at least 48 h. The digested samples were diluted 10-fold with 1% nitric acid solution containing 50 μg/L Ir as internal standard. Matrix-matched standard solutions containing Mn, Fe, Co, Cu, Zn, Ni, Ca and S were prepared in an identical manner for generation of a calibration curve. All standard and sample solutions were analyzed by inductively coupled plasma optical emission spectrometry (ICP-OES) on a Thermo iCAP PRO instrument (RF power 1250 W, with nebulizer gas flow 0.5 L/min, auxiliary gas flow 0.5 L/min, and cool gas flow 13.5 L/min argon). Elemental concentrations in each sample were calculated by comparison with the standard curve with Qtegra ISDS Software (Thermo).

### Csp *in vitro* copper binding assay

Purified Csp (500 μL, estimated at 10 μM by nanodrop) was digested in nitric acid and analyzed by ICP-OES. Protein concentrations were then accurately calculated from the quantitation of S (Csp contains 14 Cys and 4 Met residues). Purified Csp was buffer-exchanged into 20 mM Tris, pH 7.5, 150 mM NaCl to remove EDTA from the purification process using Amicon filter-concentrators (10 kDa MWCO). Aliquots (25 μM) were then titrated with increasing equivalents Cu(I) by addition of a 1:20 mixture of CuSO_4_ and ascorbate. After each addition of 1 mole equivalent Cu(I), samples were incubated for 5 minutes at room temperature. After titration with 10 and 20 mole equivalents, aliquots (500 μL) were resolved by SEC on Superdex 75 in the same buffer. Fractions were analyzed by ICP-OES as above.

### Growth curves

Bacteria were grown as described above. Suspensions were grown for 2 h, diluted to OD_600_ ∼ 0.02 (∼ 2 - 4 x 10^7^ bacteria/ml) and grown for additional 6 h. Growth was monitored hourly spectrophotometrically at OD_600_ and by plating serial culture dilutions on GC plates in triplicate for counting of colony forming units (CFU).

### Mouse antibody ELISA

Bacterial suspensions (*N. gonorrhoeae* F62 wildtype, Δ*csp*, *csp_c* and *csp_c+*) were grown as described above, centrifuged and resuspended at OD_600_ of ∼ 2 (corresponding to ∼ 2-4 x 10^9^ bacteria/ml) followed by formalin fixing as previously described [8]. ELISA plates were coated with 100 µl of bacterial suspensions diluted to obtain ∼ 1.5 x 10^8^ bacteria/ml, or with purified recombinant Csp (2 μg/ml) overnight at 4°C. Plates were then washed, blocked with 1% BSA in PBS/0.05% Tween-20 (PBS/T) for 2 h at room temperature (r.t.), and incubated with serial dilutions of pooled mouse sera (alum alone or anti-NGO1701) overnight at 4°C. The next day, plates were washed, incubated for 2h at r.t. with AP-conjugated secondary anti-mouse total IgG or IgM antibodies (Southern Biotech), followed by 1-step PNPP (p-nitrophenyl phosphate) reagent (Thermo Fisher Scientific) and spectrophotometric detection at OD_405._ Sera were tested in triplicate or quadruplicate wells. Total IgG and IgM were quantified in µg/ml using antibody reference standard curves (Southern Biotech) and a linear regression function [8].

### Metal sensitivity

Bacterial suspensions as described above were diluted to OD_600_ ∼ 0.02 (Time 0) and grown for additional 6 h in the presence of 10, 50, 100, 200 or 500 µM copper sulphate (CuSO_4_). Growth was monitored hourly spectrophotometrically at OD_600_, and by plating serial dilutions of the cultures in triplicate on GC plates for colony counting. Each growth curve was repeated at least twice. Other metals tested included cobalt chloride (CoCl_2_, 10 or 50 µM); manganese sulphate (MnSO_4_, 25 or 50 µM); nickel sulphate (NiSO_4_, 10, 50, 200, 500 or 1500 µM), zinc chloride (ZnCl_2_, 10 or 50 µM), ferric nitrate (Fe(NO_3_)_3_, 100 µM for iron-replete conditions) and Desferal (100 μM for iron-depleted conditions).

### Disc diffusion assay

Bacterial suspensions as above were diluted to OD_600_ ∼ 0.2 (2-4 x 10^8^ bacteria/ml) and plated as a lawn on GC plates using a sterile cotton swab. Paper discs (Oxoid) pre-soaked with 10 µl of CuSO_4_ solution at concentrations of 20, 50, 100 or 200 mM were placed on the lawn and plates were incubated overnight [46]. Discs soaked with sterile water were used as negative control. The next day, the zone of clearing around each disc was measured in mm. Disc diffusion experiments were repeated three times per each strain.

### Measurement of intracellular copper content

Bacterial suspensions as above were diluted to OD_600_ of ∼ 0.1 and incubated with 200 µM copper sulphate for 4 h. Bacteria were pelleted, washed with 5 ml of cold PBS, followed by another wash with PBS containing 1 mM EDTA to remove any additional residual metal. Bacteria were heat-killed at 65°C for 30 minutes and pellets were digested in 500 μL concentrated (65% v/v) nitric acid (Merck) at 70°C for at least 48 h. Once digestion was complete, the digests were centrifuged at 10,000 *g* for 10 minutes and the supernatants diluted 10-fold with 1% nitric acid solution and analyzed by ICP-OES as described above. Metal contents were normalized for small differences in biomass according to the S measurements to yield metal:sulfur ratios to enable comparisons.

### HeLa cells stimulation and invasion assay

HeLa cells (ATCC CCL-2) were grown in DMEM medium (Gibco) containing 10% FBS, 2 mM L-glutamine, 100 U/ml penicillin and 100 μg/ml streptomycin in a 5% CO_2_ incubator at 37°C. For stimulation, 10^4^ cells/well were seeded in 96-well plates and let adhere overnight. The next day, formalin-killed *N. gonorrhoeae* F62, *Δcsp*, *csp_c* and *csp_c*+ (∼ 1-2 x 10^7^ total bacteria in 100 µl) and purified Csp (10 μg/ml) were added to the wells and incubated overnight. Medium alone was used as negative control. Supernatants were collected, and secretion of IL-8 was quantified by ELISA using OptEIA ELISA kits (BD Biosciences) as specified by the manufacturer. For bacterial adherence and internalization assays, HeLa cells (5x10^4^ cells/ml) were seeded in 24-well plates as above. The next day, wells were washed with PBS, followed by addition of live *N. gonorrhoeae* F62, *Δcsp*, *csp_c* or *csp_c+* suspensions at MOI 100 (multiplicity of infection) in fresh medium without antibiotics. The co-cultures were incubated for 2 h at 37°C, after which wells were washed with PBS to remove free and non-adherent bacteria. To evaluate bacterial adherence, a set of wells was incubated with 200 μl of PBS containing 1% saponin for 10 minutes at room temperature to lyse the cells followed by vigorous pipetting. Cell lysates were serially diluted and plated on GC plates for subsequent colony counting. Invasion was evaluated by gentamicin protection assay [47]. Briefly, fresh medium containing 100 µg/ml of gentamicin (Sigma) was added to another set of wells for 1 h at 37° C to kill adherent bacteria. Wells were washed with PBS, followed by addition of fresh medium without gentamicin overnight. The next day, cells were lysed as described above to release intracellular bacteria, and the lysates were plated as above. Experiments were repeated three times.

### Serum Bactericidal Activity assay (SBA)

SBA was carried out in 96-well U-bottom plates as previously described [8, 11]. Bacterial suspensions (OD_600_ of ∼ 0.2) were serially diluted to 2–4 x 10^4^ bacteria/ml in HBSS supplemented with 0.15 mM CaCl_2_, 1 mM MgCl_2_ (HBSS^++^) and 2% BSA [11]. Bacteria were incubated with serial dilutions of heat-inactivated (56 °C for 30 min) pooled mouse sera (alum alone or anti-NGO1701) for 20 min at 37 °C. IgG/IgM-depleted normal human serum (NHS) (10%) (Pel-Freez Biologicals, Rogers, AR) was added as a source of complement, and 5 µl of bacterial suspensions were immediately plated in triplicate on GC plates (Time 0). The remaining suspensions were incubated for 30 minutes at 37 °C and aliquots were plated as above (Time 30). The next day, bacterial killing was evaluated by CFU counting. Survival was expressed as percent of CFUs at T30/T0, and bactericidal titers were defined as the reciprocal of the lowest serum dilution with ≥ 50% killing after 30 minutes. Controls included bacteria alone and bacteria incubated with NHS alone.

### Statistical Analysis

Statistical significance was examined with GraphPad Prism 10.4.1 by Ordinary one-way ANOVA with Tukey’s comparison test, by 2way ANOVA with Tukey’s multiple comparisons test or with Dunnett’s multiple comparisons test, by multiple Mann-Whitney test with Holm-Sidak correction method (p = 0.05) or by unpaired t test as indicated in the Figure legends. Differences were considered significant at a minimum *p* value of 0.05.

## Acknowledgments

The authors thank Dr. Karrera Djoko (Durham University, UK) for critical reading of the manuscript.

## Data Availability

All relevant data are within the manuscript and its Supporting information files.

## References

1. CDC. Sexually Transmitted Disease Surveillance 2023 2023. Available from: https://www.cdc.gov/sti-statistics/annual/index.html.

2. Workowski KA, Bachmann LH, Chan PA, Johnston CM, Muzny CA, Park I, et al. Sexually Transmitted Infections Treatment Guidelines, 2021. MMWR Recomm Rep. 2021;70(4):1-187. Epub 20210723. doi: 10.15585/mmwr.rr7004a1. PubMed PMID: 34292926; PubMed Central PMCID: PMCPMC8344968.

3. Unemo M, Del Rio C, Shafer WM. Antimicrobial Resistance Expressed by Neisseria gonorrhoeae: A Major Global Public Health Problem in the 21st Century. Microbiol Spectr. 2016;4(3). Epub 2016/06/24. doi: 10.1128/microbiolspec.EI10-0009-2015. PubMed PMID: 27337478; PubMed Central PMCID: PMCPMC4920088.

4. Le Van A, Rahman N, Sandy R, Dozier N, Smith HJ, Martin MJ, et al. Common Patterns and Unique Threats in Antimicrobial Resistance as Demonstrated by Global Gonococcal Surveillance. Emerg Infect Dis. 2024;30(14):62–70. doi: 10.3201/eid3014.240296. PubMed PMID: 39530861; PubMed Central PMCID: PMCPMC11559582.

5. Williams E, Seib KL, Fairley CK, Pollock GL, Hocking JS, McCarthy JS, et al. Neisseria gonorrhoeae vaccines: a contemporary overview. Clin Microbiol Rev. 2024;37(1):e0009423. Epub 20240116. doi: 10.1128/cmr.00094-23. PubMed PMID: 38226640; PubMed Central PMCID: PMCPMC10938898.

6. Gulati S, Shaughnessy J, Ram S, Rice PA. Targeting Lipooligosaccharide (LOS) for a Gonococcal Vaccine. Front Immunol. 2019;10:321. Epub 2019/03/16. doi: 10.3389/fimmu.2019.00321. PubMed PMID: 30873172; PubMed Central PMCID: PMCPMC6400993.

7. Gulati S, Pennington MW, Czerwinski A, Carter D, Zheng B, Nowak NA, et al. Preclinical Efficacy of a Lipooligosaccharide Peptide Mimic Candidate Gonococcal Vaccine. mBio. 2019;10(6). Epub 2019/11/07. doi: 10.1128/mBio.02552-19. PubMed PMID: 31690678; PubMed Central PMCID: PMCPMC6831779.

8. Zhu T, McClure R, Harrison OB, Genco C, Massari P. Integrated Bioinformatic Analyses and Immune Characterization of New Neisseria gonorrhoeae Vaccine Antigens Expressed during Natural Mucosal Infection. Vaccines (Basel). 2019;7(4). Epub 20191017. doi: 10.3390/vaccines7040153. PubMed PMID: 31627489; PubMed Central PMCID: PMCPMC6963464.

9. McClure R, Nudel K, Massari P, Tjaden B, Su X, Rice PA, et al. The Gonococcal Transcriptome during Infection of the Lower Genital Tract in Women. PLoS One. 2015;10(8):e0133982. Epub 20150805. doi: 10.1371/journal.pone.0133982. PubMed PMID: 26244506; PubMed Central PMCID: PMCPMC4526530.

10. Nudel K, McClure R, Moreau M, Briars E, Abrams AJ, Tjaden B, et al. Transcriptome Analysis of Neisseria gonorrhoeae during Natural Infection Reveals Differential Expression of Antibiotic Resistance Determinants between Men and Women. mSphere. 2018;3(3). Epub 2018/06/29. doi: 10.1128/mSphereDirect.00312-18. PubMed PMID: 29950382; PubMed Central PMCID: PMCPMC6021601.

11. Roe SK, Felter B, Zheng B, Ram S, Wetzler LM, Garges E, et al. In Vitro Pre-Clinical Evaluation of a Gonococcal Trivalent Candidate Vaccine Identified by Transcriptomics. Vaccines (Basel). 2023;11(12). Epub 20231213. doi: 10.3390/vaccines11121846. PubMed PMID: 38140249; PubMed Central PMCID: PMCPMC10747275.

12. Vita N, Platsaki S, Basle A, Allen SJ, Paterson NG, Crombie AT, et al. A four-helix bundle stores copper for methane oxidation. Nature. 2015;525(7567):140–3. Epub 2015/08/27. doi: 10.1038/nature14854. PubMed PMID: 26308900; PubMed Central PMCID: PMCPMC4561512.

13. Vita N, Landolfi G, Basle A, Platsaki S, Lee J, Waldron KJ, et al. Bacterial cytosolic proteins with a high capacity for Cu(I) that protect against copper toxicity. Sci Rep. 2016;6:39065. Epub 20161219. doi: 10.1038/srep39065. PubMed PMID: 27991525; PubMed Central PMCID: PMCPMC5171941.

14. Focarelli F, Giachino A, Waldron KJ. Copper microenvironments in the human body define patterns of copper adaptation in pathogenic bacteria. PLoS Pathog. 2022;18(7):e1010617. Epub 20220721. doi: 10.1371/journal.ppat.1010617. PubMed PMID: 35862345; PubMed Central PMCID: PMCPMC9302775.

15. Jolley KA, Bray JE, Maiden MCJ. Open-access bacterial population genomics: BIGSdb software, the PubMLST.org website and their applications. Wellcome Open Res. 2018;3:124. Epub 20180924. doi: 10.12688/wellcomeopenres.14826.1. PubMed PMID: 30345391; PubMed Central PMCID: PMCPMC6192448.

16. Archie Howell SC, Karrera Y. Djoko. Copper homeostasis in Streptococcus and Neisseria: Known knowns and unknown knowns. Advances in Microbial Physiology. 2025. doi: 10.1016/bs.ampbs.2024.11.001.

17. Stohl EA, Lenz JD, Dillard JP, Seifert HS. The Gonococcal NlpD Protein Facilitates Cell Separation by Activating Peptidoglycan Cleavage by AmiC. J Bacteriol. 2015;198(4):615–22. Epub 20151116. doi: 10.1128/JB.00540-15. PubMed PMID: 26574512; PubMed Central PMCID: PMCPMC4751805.

18. Lu P, Wang S, Lu Y, Neculai D, Sun Q, van der Veen S. A Subpopulation of Intracellular Neisseria gonorrhoeae Escapes Autophagy-Mediated Killing Inside Epithelial Cells. J Infect Dis. 2019;219(1):133–44. doi: 10.1093/infdis/jiy237. PubMed PMID: 29688440.

19. Inacio AS, Nunes A, Milho C, Mota LJ, Borrego MJ, Gomes JP, et al. In Vitro Activity of Quaternary Ammonium Surfactants against Streptococcal, Chlamydial, and Gonococcal Infective Agents. Antimicrob Agents Chemother. 2016;60(6):3323–32. Epub 20160523. doi: 10.1128/AAC.00166-16. PubMed PMID: 26976875; PubMed Central PMCID: PMCPMC4879390.

20. Mavrogiorgos N, Mekasha S, Yang Y, Kelliher MA, Ingalls RR. Activation of NOD receptors by Neisseria gonorrhoeae modulates the innate immune response. Innate Immun. 2014;20(4):377–89. Epub 20130724. doi: 10.1177/1753425913493453. PubMed PMID: 23884094; PubMed Central PMCID: PMCPMC3880408.

21. Jones A, Jonsson AB, Aro H. Neisseria gonorrhoeae infection causes a G1 arrest in human epithelial cells. FASEB J. 2007;21(2):345–55. Epub 20061208. doi: 10.1096/fj.06-6675com. PubMed PMID: 17158783.

22. Petousis-Harris H, Paynter J, Morgan J, Saxton P, McArdle B, Goodyear-Smith F, et al. Effectiveness of a group B outer membrane vesicle meningococcal vaccine against gonorrhoea in New Zealand: a retrospective case-control study. Lancet. 2017;390(10102):1603–10. Epub 2017/07/15. doi: 10.1016/S0140-6736(17)31449-6. PubMed PMID: 28705462.

23. Abara WE, Kirkcaldy RD, Bernstein KT, Galloway E, Learner ER. Effectiveness of MenB-4C Vaccine Against Gonorrhea: A Systematic Review and Meta-analysis. J Infect Dis. 2025;231(1):61–70. doi: 10.1093/infdis/jiae383. PubMed PMID: 39082700; PubMed Central PMCID: PMCPMC11782638.

24. Leduc I, Connolly KL, Begum A, Underwood K, Darnell S, Shafer WM, et al. The serogroup B meningococcal outer membrane vesicle-based vaccine 4CMenB induces cross-species protection against Neisseria gonorrhoeae. PLoS Pathog. 2020;16(12):e1008602. Epub 2020/12/09. doi: 10.1371/journal.ppat.1008602. PubMed PMID: 33290434; PubMed Central PMCID: PMCPMC7748408.

25. Viviani V, Fantoni A, Tomei S, Marchi S, Luzzi E, Bodini M, et al. OpcA and PorB are novel bactericidal antigens of the 4CMenB vaccine in mice and humans. NPJ Vaccines. 2023;8(1):54. Epub 20230412. doi: 10.1038/s41541-023-00651-9. PubMed PMID: 37045859; PubMed Central PMCID: PMCPMC10097807.

26. Matthias KA, Connolly KL, Begum AA, Jerse AE, Macintyre AN, Sempowski GD, et al. Meningococcal Detoxified Outer Membrane Vesicle Vaccines Enhance Gonococcal Clearance in a Murine Infection Model. J Infect Dis. 2022;225(4):650–60. doi: 10.1093/infdis/jiab450. PubMed PMID: 34498079; PubMed Central PMCID: PMCPMC8844591.

27. Liu Y, Hammer LA, Daamen J, Stork M, Egilmez NK, Russell MW. Microencapsulated IL-12 Drives Genital Tract Immune Responses to Intranasal Gonococcal Outer Membrane Vesicle Vaccine and Induces Resistance to Vaginal Infection with Diverse Strains of Neisseria gonorrhoeae. mSphere. 2023;8(1):e0038822. Epub 2022/12/21. doi: 10.1128/msphere.00388-22. PubMed PMID: 36537786; PubMed Central PMCID: PMCPMC9942569.

28. Dhital S, Deo P, Stuart I, Huang C, Zavan L, Han ML, et al. Characterization of outer membrane vesicles released by clinical isolates of Neisseria gonorrhoeae. Proteomics. 2024;24(11):e2300087. Epub 20231207. doi: 10.1002/pmic.202300087. PubMed PMID: 38059892.

29. Marjuki H, Topaz N, Joseph SJ, Gernert KM, Kersh EN, Antimicrobial-Resistant Neisseria gonorrhoeae Working G, et al. Genetic Similarity of Gonococcal Homologs to Meningococcal Outer Membrane Proteins of Serogroup B Vaccine. mBio. 2019;10(5). Epub 2019/09/12. doi: 10.1128/mBio.01668-19. PubMed PMID: 31506309; PubMed Central PMCID: PMCPMC6737241.

30. Matthias KA, Reveille A, Dhara K, Lyle CS, Natuk RJ, Bonk B, et al. Development and validation of a standardized human complement serum bactericidal activity assay to measure functional antibody responses to Neisseria gonorrhoeae. Vaccine. 2025;43(Pt 2):126508. Epub 20241115. doi: 10.1016/j.vaccine.2024.126508. PubMed PMID: 39549368.

31. Straw ML, Chaplin AK, Hough MA, Paps J, Bavro VN, Wilson MT, et al. A cytosolic copper storage protein provides a second level of copper tolerance in Streptomyces lividans. Metallomics. 2018;10(1):180–93. doi: 10.1039/c7mt00299h. PubMed PMID: 29292456.

32. Lee J, Dalton RA, Dennison C. Copper delivery to an endospore coat protein of Bacillus subtilis. Front Cell Dev Biol. 2022;10:916114. Epub 20220905. doi: 10.3389/fcell.2022.916114. PubMed PMID: 36133923; PubMed Central PMCID: PMCPMC9484137.

33. Branch AH, Stoudenmire JL, Seib KL, Cornelissen CN. Acclimation to Nutritional Immunity and Metal Intoxication Requires Zinc, Manganese, and Copper Homeostasis in the Pathogenic Neisseriae. Front Cell Infect Microbiol. 2022;12:909888. Epub 2022/07/19. doi: 10.3389/fcimb.2022.909888. PubMed PMID: 35846739; PubMed Central PMCID: PMCPMC9280163.

34. Liyayi IK, Forehand AL, Ray JC, Criss AK. Metal piracy by Neisseria gonorrhoeae to overcome human nutritional immunity. PLoS Pathog. 2023;19(2):e1011091. Epub 20230202. doi: 10.1371/journal.ppat.1011091. PubMed PMID: 36730177; PubMed Central PMCID: PMCPMC9894411.

35. Baarda BI, Zielke RA, Jerse AE, Sikora AE. Lipid-Modified Azurin of Neisseria gonorrhoeae Is Not Surface Exposed and Does Not Interact With the Nitrite Reductase AniA. Front Microbiol. 2018;9:2915. Epub 2018/12/13. doi: 10.3389/fmicb.2018.02915. PubMed PMID: 30538694; PubMed Central PMCID: PMCPMC6277709.

36. Barreiro DS, Oliveira RNS, Pauleta SR. Biochemical Characterization of the Copper Nitrite Reductase from Neisseria gonorrhoeae. Biomolecules. 2023;13(8). Epub 20230804. doi: 10.3390/biom13081215. PubMed PMID: 37627281; PubMed Central PMCID: PMCPMC10452240.

37. Jen FE, Djoko KY, Bent SJ, Day CJ, McEwan AG, Jennings MP. A genetic screen reveals a periplasmic copper chaperone required for nitrite reductase activity in pathogenic Neisseria. FASEB J. 2015;29(9):3828–38. Epub 2015/06/03. doi: 10.1096/fj.15-270751. PubMed PMID: 26031293.

38. McClure R, Sunkavalli A, Balzano PM, Massari P, Cho C, Nauseef WM, et al. Global Network Analysis of Neisseria gonorrhoeae Identifies Coordination between Pathways, Processes, and Regulators Expressed during Human Infection. mSystems. 2020;5(1). Epub 20200204. doi: 10.1128/mSystems.00729-19. PubMed PMID: 32019834; PubMed Central PMCID: PMCPMC7002116.

39. Costa-Lourenco APR, Su X, Le W, Yang Z, Patts GJ, Massari P, et al. Epidemiological and Clinical Observations of Gonococcal Infections in Women and Prevention Strategies. Vaccines (Basel). 2021;9(4). Epub 2021/05/01. doi: 10.3390/vaccines9040327. PubMed PMID: 33915835; PubMed Central PMCID: PMCPMC8066387.

40. Madeira F, Madhusoodanan N, Lee J, Eusebi A, Niewielska A, Tivey ARN, et al. The EMBL-EBI Job Dispatcher sequence analysis tools framework in 2024. Nucleic Acids Res. 2024;52(W1):W521-W5. doi: 10.1093/nar/gkae241. PubMed PMID: 38597606; PubMed Central PMCID: PMCPMC11223882.

41. McGuffin LJ, Adiyaman R, Maghrabi AHA, Shuid AN, Brackenridge DA, Nealon JO, et al. IntFOLD: an integrated web resource for high performance protein structure and function prediction. Nucleic Acids Res. 2019;47(W1):W408–W13. doi: 10.1093/nar/gkz322. PubMed PMID: 31045208; PubMed Central PMCID: PMCPMC6602432.

42. Baek M, Park T, Heo L, Park C, Seok C. GalaxyHomomer: a web server for protein homo-oligomer structure prediction from a monomer sequence or structure. Nucleic Acids Res. 2017;45(W1):W320–W4. doi: 10.1093/nar/gkx246. PubMed PMID: 28387820; PubMed Central PMCID: PMCPMC5570155.

43. The PyMOL Molecular Graphics System VS, LLC. Available from: http://www.pymol.org/pymol.

44. Dillard JP. Genetic Manipulation of Neisseria gonorrhoeae. Curr Protoc Microbiol. 2011;Chapter 4:Unit4A 2. doi: 10.1002/9780471729259.mc04a02s23. PubMed PMID: 22045584; PubMed Central PMCID: PMCPMC4549065.

45. Skaar EP, Lazio MP, Seifert HS. Roles of the recJ and recN genes in homologous recombination and DNA repair pathways of Neisseria gonorrhoeae. J Bacteriol. 2002;184(4):919–27. doi: 10.1128/jb.184.4.919-927.2002. PubMed PMID: 11807051; PubMed Central PMCID: PMCPMC134828.

46. Djoko KY, Franiek JA, Edwards JL, Falsetta ML, Kidd SP, Potter AJ, et al. Phenotypic characterization of a copA mutant of Neisseria gonorrhoeae identifies a link between copper and nitrosative stress. Infect Immun. 2012;80(3):1065–71. Epub 2011/12/21. doi: 10.1128/IAI.06163-11. PubMed PMID: 22184419; PubMed Central PMCID: PMCPMC3294635.

47. Almonacid-Mendoza HL, Christodoulides M. Basic Methods for Examining Neisseria gonorrhoeae Interactions with Host Cells In Vitro. Methods Mol Biol. 2019;1997:281–99. doi: 10.1007/978-1-4939-9496-0_17. PubMed PMID: 31119630.

