## Supplemental Materials for "The gonococcal vaccine candidate antigen NGO1701 is a *N. gonorrhoeae* periplasmic copper storage protein"

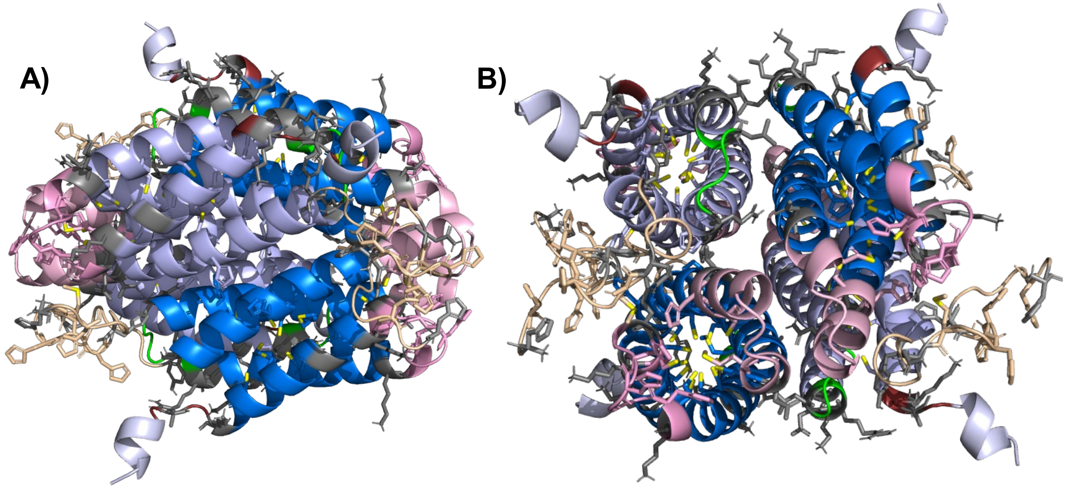


**Supplemental Figure 1**

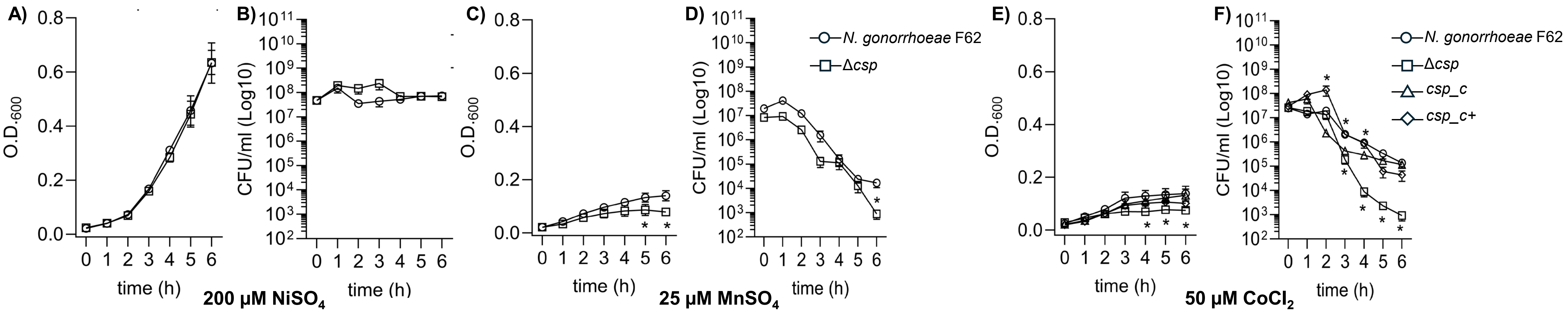


**Supplemental Figure 2**

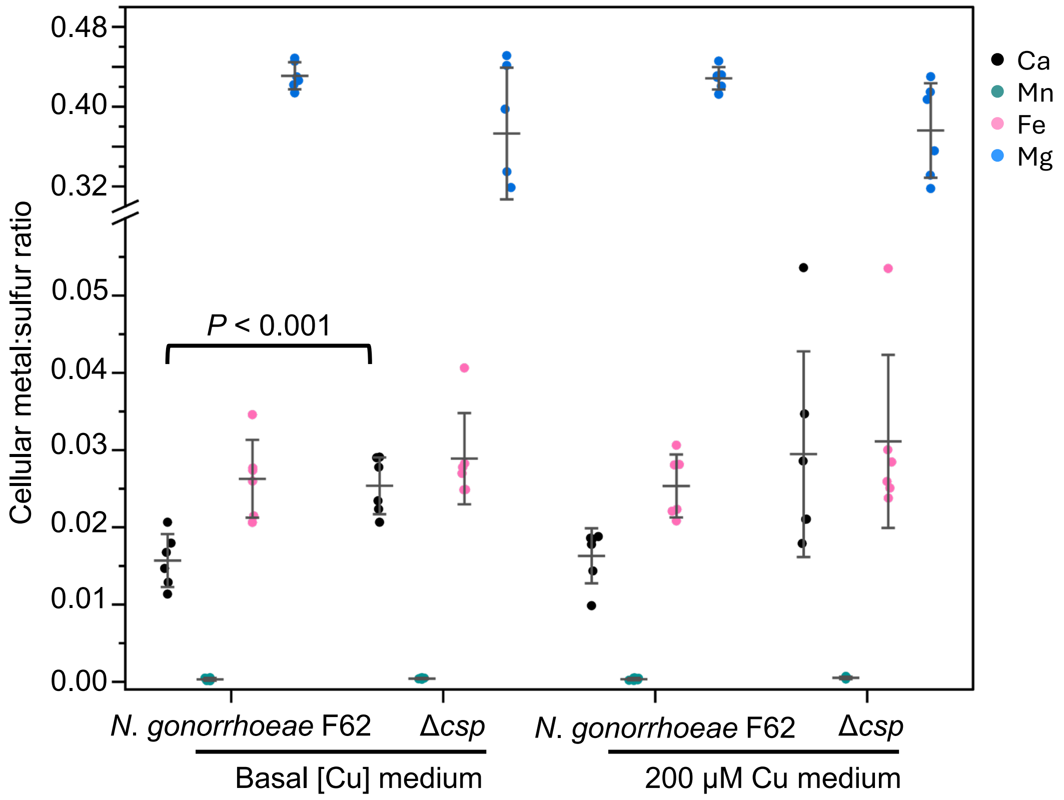


**Supplemental Figure 3**

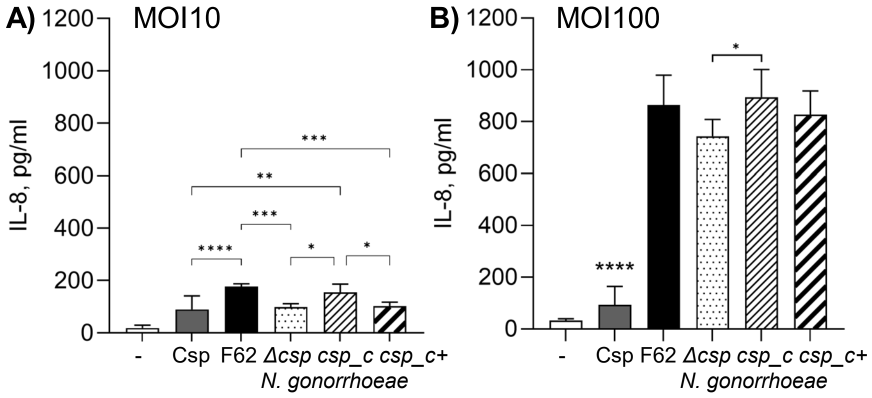


**Supplemental Figure 4**

**Supplemental Table 1. List of primers and sequences**

| Primer name | Sequence (5'-3') |
|---|---|
| *ngo1701* upstream_F | GGCCAAAGGCATGAAGGGAATAAAGGG |
| *ngo1701* upstream_R | CGGATGGAACCAGGCTGCATATCGTGATGATGGTGGTAG |
| *ngo1701* downstream_F | GCAAAGAAGCTTCGAGTCCTGCCTCGACTGTATCAAAGAATG |
| *ngo1701* downstream_R | TTGCCCGATGTGGCGATTGCTAAAG |
| pGCC4 Kan_F | CATTTCGAATTCCAGAGTCCCGCTCAGAAGAAC |
| pGCC4 Kan_R | ATGGACAAGCTTCGAACCGGAATTGCCAGCTGG |
| *ngo1701*_F check | TGCAGCAGATGATGAAGATGTTC |
| pGCC4 mid_Kan | TACGCTTGATCCGGCTACCT |
| *lctP*_F | CTGTCCGTGACCCTGATTCTGGC |
| *acpC*_R | GATGCCGAGGTTGACTTTTT |
| *lctP*_F and *lacIq*_R | CCACCCTGAATTGACTCTCTTCCGG |

**Supplemental Figure 1. Structure model of the Csp tetramer. A)** Side view of a cartoon model of Csp based on the Csp1 structural model created with GalaxyHomomer (one pair of symmetry-related monomers shown in dark blue and the other pair in light blue). **B)** Front view (90° rotation relative to A). The colored regions illustrate the predicted linear epitopes (LE) 1-5 and conformational epitopes (CE) 1-3 [11], located on the outside of the tetrameric structure, validating the model and confirming this tetrameric form as the likely structure in solution.

**Supplemental Figure 2. *N. gonorrhoeae* growth and metal toxicity.** $OD_{600}$ values (average ± SEM) from three independent growth curves in the presence of **A)** 200 μM nickel sulfate ($NiSO_4$), **C)** 25 μM manganese sulfate ($MnSO_4$) or **E)** 50 μM cobalt chloride ($CoCl_2$). * $p \leq 0.05$ vs no treatment by 2-way ANOVA with Dunnett's multiple comparisons test. **B-D-F)** CFUs/ml (average ± SEM) as above. Statistical significance was determined by multiple Mann-Whitney test with Holm-Sidak correction set for p = 0.05 vs no treatment for each strain, indicated by *. *N. gonorrhoeae* F62 wildtype (circles), Δ*csp* (squares), *csp_c* (triangles) and *csp_c*+ (diamonds).

**Supplemental Figure 3. Accumulation of other metals in *N. gonorrhoeae* Δ*csp* is not affected by Cu.** *N. gonorrhoeae* F62 wildtype and Δ*csp* were grown for 4 h in GBC without added copper (basal [Cu] medium) or in the presence of 200 μM copper sulphate. After removing residual surface-adsorbed metal, bacterial pellets were digested and analyzed for Ca (black), Mn (teal), Fe (pink), Mg (blue) and S by ICP-OES. Metal content was normalized to S content to account for differences in biomass. *P* represents the results of an unpaired *t* test.

**Supplemental Figure 4. IL-8 secretion by HeLa cells.** Cells were incubated with purified Csp (10 μg/ml) (gray bars), *N. gonorrhoeae* F62 wildtype (black bars), Δ*csp* (dotted bars), *csp_c* (thin striped bars) or *csp_c*+ (thick striped bars) at **A)** MOI 10 or **B)** MOI 100 for 18 h. IL-8 secretion was measured in cell culture supernatants by ELISA and expressed as pg/ml ±

SEM from triplicate wells. * $p < 0.05$, **, $p < 0.005$, ***, $p < 0.0005$ and ****, $p < 0.0001$ by Ordinary one way ANOVA with Tukey's multiple comparison test.
